# A Comprehensive and Robust Multiplex-DIA Workflow Profiles Protein Turnover Regulations Associated with Cisplatin Resistance

**DOI:** 10.1101/2024.10.28.620709

**Authors:** Barbora Salovska, Wenxue Li, Oliver M. Bernhardt, Pierre-Luc Germain, Tejas Gandhi, Lukas Reiter, Yansheng Liu

## Abstract

Measuring protein turnover is essential for understanding cellular biological processes and advancing drug discovery. The multiplex DIA mass spectrometry (DIA-MS) approach, combined with dynamic SILAC labeling (pulse-SILAC, or pSILAC), has proven to be a reliable method for analyzing protein turnover and degradation kinetics. Previous multiplex DIA-MS workflows have employed various strategies, including leveraging the highest isotopic labeling channels of peptides to enhance the detection of isotopic MS signal pairs or clusters. In this study, we introduce an improved and robust workflow that integrates a novel machine learning strategy and channel-specific statistical filtering, enabling dynamic adaptation to systematic or temporal variations in channel ratios. This allows comprehensive profiling of protein turnover throughout the pSILAC experiment without relying solely on the highest channel signals. Additionally, we developed ***KdeggeR***, a data processing and analysis package optimized for pSILAC-DIA experiments, which estimates and visualizes peptide and protein degradation rates and dynamic profiles. Our integrative workflow was benchmarked on both 2-channel and 3-channel standard DIA datasets and workflow to an aneuploid cancer cell model before and after cisplatin resistance development demonstrated a strong negative correlation between transcript regulation and protein degradation for major protein complex subunits. We also identified specific protein turnover signatures associated with cisplatin resistance.

## Introduction

Protein turnover, the balance between synthesis and degradation, is a fundamental process that regulates cellular homeostasis, adaptation, and response to environmental stimuli. It plays a crucial role in a variety of biological processes, including cell growth, differentiation, and apoptosis, and is a critical factor in understanding disease progression and therapeutic responses. In cancer biology, for example, altered protein turnover rates are often linked to genomic instability, such as aneuploidy, and resistance to chemotherapeutic agents like cisplatin ^1,2^. Understanding the dynamics of protein turnover in these contexts is essential for identifying potential therapeutic targets and biomarkers.

Mass spectrometry (MS)-based approaches have become a key tool for studying protein turnover, with data-independent acquisition (DIA) MS being one of the most robust and reproducible techniques ^3,4^. The multiplex DIA-MS approach, when combined with dynamic stable isotope labeling by amino acids in cell culture (pulse-SILAC, or pSILAC)) ^5–8^, allows for multi-time-point measurements and the precise quantification of protein-specific turnover rates across diverse biological conditions ^9–14^. Its ability to profile large numbers of peptides and proteomes reproducibly makes DIA-MS particularly well-suited for complex experimental designs, such as time-course experiments often used in dynamic SILAC design.

Recent advancements in MS acquisition strategies have further enhanced the power and throughput of DIA-MS workflows. Techniques such as BoxCarmax ^15^, which involves small isolation windows in combination with multiple runs, and instruments like Astral ^16^ and timsTOF, which support small isolation windows directly, have significantly improved the peptide detection and quantification of heavy (H) and light (L) ratios. These advancements increase the precision and depth of protein turnover analysis.

A particular challenge that pSILAC experiments face is that the intensity of the channels mirrors the protein turnover rate which can lead to a near absence of one of the channels for a particular protein (or peptide) in the early and very late time points. Previous strategies have relied on focusing on specific channels: We previously presented an Inverted Spike-In Workflow (ISW) ^10^ which utilizes only the light channel for scoring and signal detection in pSILAC-DIA data, increasing the number of H/L pairs being measured by ca. 30% in early pSILAC labeling time points. However, ISW is not ideal for the late pSILAC labeling time points and other multiplex DIA-MS experiments in which the labeled peptide signals are often higher than the light ones depending on the specific experimental condition and individual proteins. On the other hand, strategies such as plex-DIA ^17^ or mDIA ^18^ utilize DIA-NN ^19^ or RefQuant ^18^ software tools to target the highest isotopic channel or the reference channel for improving peptide detection. These approaches, while effective, still leave room for optimization, in e.g., leveraging all available isotopic channels and peptide transitions for more comprehensive protein turnover quantification.

Herein, we introduce a novel approach that incorporates machine learning and channel-specific statistical filtering into the peptide detection process in multiplex DIA datasets. Our method dynamically adapts to systematic changes in isotopic channel ratios, ensuring that all channels are effectively utilized without the need of selecting the highest signals. We extensively assessed this improvement and found that, together with a following data processing tool, the accuracy and robustness of protein turnover measurements in complex pSILAC datasets were significantly enhanced.

To apply our enhanced workflow, we focused on an aneuploidy cancer cell model. Aneuploidy, characterized by an abnormal number of chromosomes, alters the protein homeostasis landscape, leading to unique turnover profiles that may contribute to the development of resistance to drugs. Our pSILAC-DIA measurement and workflow applied in the aneuploidy ovarian cells divergent for cisplatin resistance uncovered key turnover signatures and regulations that potentially drive resistance mechanisms, providing new insights into potential therapeutic strategies.

## Results

### Overview of a Robust Workflow for Multiplex-DIA MS Data Analysis Enabling Large-Scale Protein Turnover Quantification and Comparative Studies

Multiplex-DIA-MS, combined with pSILAC enables precise quantification of protein turnover across multiple conditions by measuring protein dynamics at different time points (**Figure 1, Upper left**). However, effectively detecting and quantifying MS signals in multiplex pSILAC-DIA datasets in which the heavy signal might be low abundant in the early labeling time points remains challenging. To address this challenge, we previously introduced the “Inverted Spike-In workflow” (ISW) ^10^. In that work, we firstly relied on an extensive hybrid library generated using label-free and multiplexed samples, both DDA-MS and DIA-MS. This library was then used to perform a targeted extraction of the multiplexed DIA-MS raw files, employing the ISW workflow, **Figure 1, Lower left**). In ISW, the peak picking and scoring are based exclusively on the “light” precursors, which we demonstrated as advantageous in samples with a low relative abundance of “heavy” signals such as extreme H/L dilutions ^15^ or early labeling time points of a pSILAC experiment ^10^.

**Figure 1:**
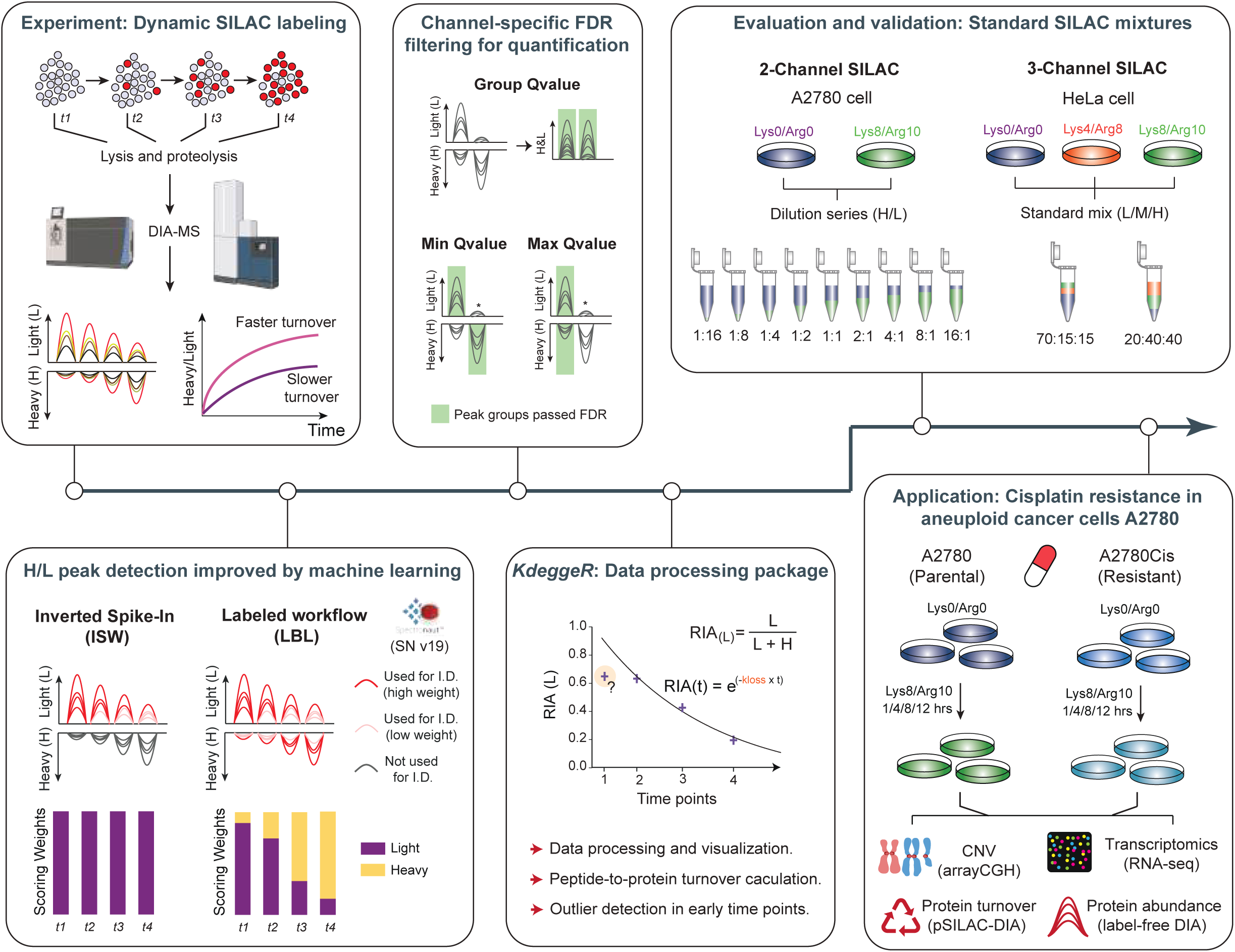
A robust workflow for multiplex-DIA MS data analysis. ***Upper left:*** Protein turnover analysis on a large scale using dynamic stable isotope labeling by amino acids in cell culture (pSILAC), combined with highly robust and reproducible multiplex data-independent acquisition (DIA) mass spectrometry (MS), enables the quantification of thousands of protein turnover rates and facilitates quantitative comparisons between multiple conditions. Datasets from two MS platforms (Orbitrap Fusion Lumos and timsTOF Ultra) were processed. ***Lower left:*** In the previously reported inverted spike-in workflow (ISW), peak picking and scoring relied solely on the light channel. In the labeled (“LBL”) workflow, XIC peak picking, elution group scoring, and “Group Qvalue” calculation are performed across all channels (n = 2, 3, …, n) in a combined fashion, facilitated by improved machine learning. ***Upper middle:*** In addition to the Group Qvalue, our Spectronaut v19 (SN19) solution now offers channel-specific Qvalue filtering options for more stringent quantification data filtering. In the “Min Qvalue” option, at least one channel needs to be independently identified (Q value < 0.01), while in the “Max Qvalue” option, all channels must be independently identified to accept the entire elution group. ***Lower middle:*** As part of the workflow, we provide an R package named ***KdeggeR*** for the analysis of pulse SILAC DIA-MS data from various raw data processing software, including data formatting, data filtering and QC, the calculation of precursor-, peptide-, and protein-level protein turnover rates (Kloss), subsequent protein degradation rate (*k*_deg_) transformation, comparative data analysis, and data visualization. ***Upper right:*** We evaluated the multichannel analysis implemented (e.g., in Spectronaut v19) using 2-channel standard dilution samples and 3-channel standard datasets. Both datasets were acquired previously and are publicly available (Salovska et al., 2021; Bortecen et al., 2024). ***Lower right:*** We demonstrated the feasibility of the entire workflow by applying it to the study of protein turnover regulation in a cisplatin resistance model of the highly aneuploid ovarian cancer A2780 and integrated the data with other omic layers. This application highlighted the importance of studying protein turnover to derive biological insights into complex phenomena such as the cancer drug resistance phenotype.

However, recent rapid advancements in library-free DIA-MS data analysis in software such as the directDIA algorithm in Spectronaut ^20^ and other software tools like DIA-NN ^19^, or FragPipe ^21^, driven by machine learning and deep learning techniques, have essentially eliminated the need to generate extensive project-specific spectral libraries for routine peptide identifications in proteomics, significantly streamlining DIA-MS data analysis. Additionally, we reason that performing the XIC peak picking and elution group scoring using the information across all channels (n = 2, 3, …, n) may enable a more dynamic scoring of increasingly complex labeling experiments, accommodating a wider range of labeled/unlabeled ratios over the entire experiment and additional labeling channels (see Introduction).

Leveraging the directDIA algorithm and improved machine learning, we optimized and evaluated a library-free “Labeled” workflow (LBL), which is available in Spectronaut v19 (**Figure 1 – Lower left**). Notably, during the targeted peak extraction, this workflow performs the XIC peak picking across all channels in a combined fashion. Moreover, in the elution group scoring, all channels, along with their specific and cross-channel scores, are considered collectively in the machine learning process, leading to an estimation of a “Group Q-value”. Together, these allow for dynamic adaptation to systematic changes in channel ratios per sample, leading to the optimal scoring weights of the labeled and unlabeled peptide transitions consistent with the SILAC ratios in real pSILAC experiments (**Figure 1 – Upper Middle**). Additionally, labeled or unlabeled channels can be also scored independently based on channel-specific metrics, supporting the determination of channel-specific Q-values, which is newly possible with Spectronaut v19. This further enables channel-specific FDR filtering of the quantification results. In the “Min Q-value” option, at least one channel needs to be independently identified (Q value < 0.01), while in the “Max Q-value” option, all channels must be independently identified to accept the entire elution group for a given peptide (**Figure 1 – Upper Middle**). To evaluate the sensitivity, quantification precision, and accuracy, we applied these integrative data procession steps to a 2-channel dilution standard dataset ^15^ and a 3-channel dataset ^22^ (**Figure 1 – Upper Right, see Methods**).

To facilitate downstream analysis of the pSILAC data, we herein also present an open-source R package, ***KdeggeR***, which aims to streamline the processing of pulse SILAC DIA-MS data. ***KdeggeR*** performs data formatting, quality control, calculation of peptide and protein turnover (*k*_loss_), degradation rates (*k*_deg_), comparative analysis, and visualization (**Figure 1, Lower Middle, see Methods**). The package supports data from various raw data processing software, making it a versatile tool for diverse proteomics workflows. Finally, we applied the entire workflow to a biological investigation on protein turnover regulation in a cisplatin-resistant ovarian cancer model (A2780 and A2780Cis cell lines). By integrating the multiplex-DIA data with other omics, we gained novel insights into the mechanisms underlying drug resistance in this highly aneuploid cancer model (**Figure 1, Lower Right**).

### Improved Identification of Multiplex-DIA-MS Datasets Through Machine Learning-Guided Dynamic Selection of Isotopic Labeling Features

To assess the effectiveness of the LBL and compare it to the ISW, we analyzed the A2780 standard 2-channel SILAC dilution series (H:L: 1:16, 1:8, …, to 8:1,16:1) as the first benchmarking dataset ^15^. We found that, in a library-free analysis of a 1:1 sample, the LBL led to the identification of 142,363 precursors and 7,785 protein groups (**Figure 2A, Figure S1A**), which was 3.8% and 12% more than the ISW result. Strikingly, this is 147.6% and 35.3% more precursors and protein groups than we reported previously in the same samples analyzed using ISW and an extensive, project-specific hybrid library (188,886 peptide precursors corresponding to 7,457 proteins) in Spectronaut v13 ^15^, demonstrating the improved software performance especially the deep learning features included in recent DIA data analysis software tools ^19–21^. Furthermore, in the dilution series of A2780, the LBL outperformed the ISW by identifying more precursors and proteins across mixing conditions (**Figure 2B, Figure S1B**). In the light-dominant samples, the LBL identified slightly more features (about 10% more precursors and 5% more protein groups), but dramatically overperformed ISW in the heavy-dominant samples, as expected. The LBL workflow successfully reached the dynamic assignment of the scoring weight to both channels (**Figure 2C**).

**Figure 2:**
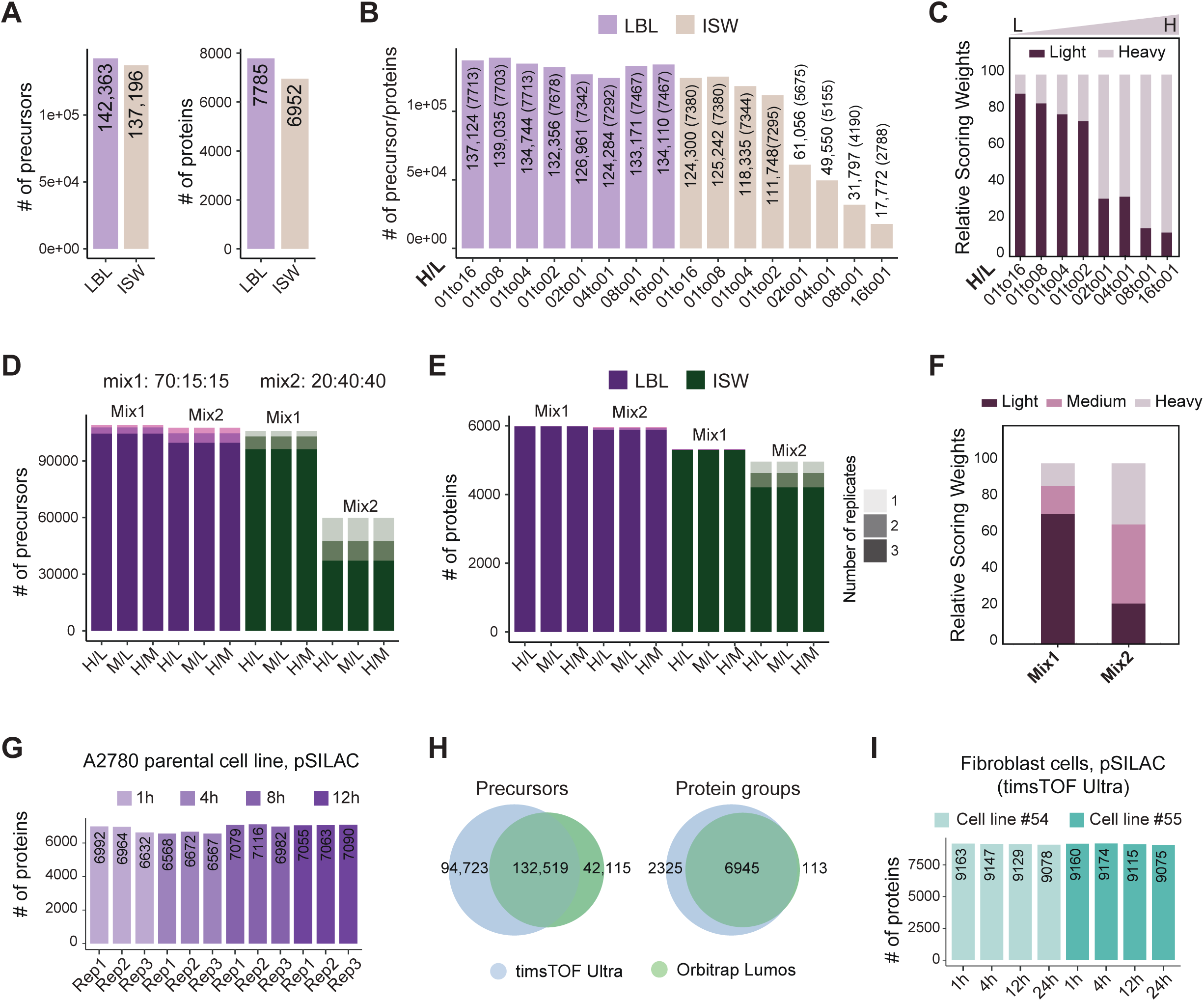
Improved identification of multiplex DIA-MS datasets using machine learning to dynamically select isotopic labeling features. **(A)** Improved identification using the Labeled workflow implemented in SN19 in the H/L = 1 sample of A2780; the numbers of identified IDs at the precursor (left) and protein (right) levels are shown. **(B)** The Labeled workflow outperformed the inverted spike-in workflow in the A2780 dilution series analysis; the numbers of identified IDs at the precursor and protein (in brackets) levels are shown. **(C)** Scoring weight histogram from the 2-channel A2780 dilution series experiment. **(D-E)** Number of precursors **(D)** and proteins **(E)** identified in the 3-channel HeLa standard sample experiment; the numbers of IDs identified in an experimental replicate are shown. **(F)** Scoring weight histogram from the 3-channel HeLa standard sample experiment. The bars represent the averaged weights per condition/mix. **(G)** Protein-level identification in the pulse SILAC experiment in the A2780 cell line. **(H)** Precursor- and protein-level comparison of identifications between samples measured using the timsTOF Ultra and Orbitrap Fusion Lumos platforms. **(I)** Protein-level identification in a pSILAC experiment.

As the second benchmarking dataset, we leveraged a public dataset of HeLa cells with 3 SILAC labeling states (Light, Medium, and Heavy) ^22^. This dataset consisted of two different compositions, i.e., **mix1** (light-dominant, H:M:L = 15:15:70) and **mix2** (medium&heavy-dominant, H:M:L = 40:40:20). In both mixes, we found that the LBL identified more precursors and proteins, leading to a greater number of pairwise ratios between the three channels (**Figure 2D, 2E, Figures S1C**). Similar to the 2-channel result, LBL yielded a slight improvement in the light-dominant **mix1** (5.3% and 12.7% more precursors and proteins, respectively), but a more dramatic improvement in **mix2** in which light peptides only account for 20% (115.9% and 28.9% more precursors and proteins, respectively). The number of missing values across the replicates was extremely low in the results based on the LBL workflow, especially at the protein level (**Figure 2E**). The scoring weight histogram of this experiment further compellingly validated the LBL algorithm (**Figure 2F**).

Next, to showcase the practical application of the LBL workflow in a real pSILAC experiment, we analyzed the third dataset obtained from the A2780 cell line (the parental line). LBL consistently identified 6,900 proteins on average, covering four time points and three experimental replicates (**Figure 2G**). Furthermore, to validate the general applicability of LBL, we analyzed a pSILAC experiment performed in two fibroblast cell lines, for which we acquired the datasets using two independent LC-MS platforms, Orbitrap Fusion Lumos and timsTOF Ultra (**Figure 2H, see Methods**). Impressively, with 2.5 times shortened gradient and less than 10% sample amount (130 ng vs 1.5 µg), using the timsTOF Ultra platform we identified in total 227,242 precursors and 9,270 protein groups (30.1% and 31.2% more than using Lumos, respectively, **Figure 2H**), and a consistent identification of 9,130 proteins on average (**Figure 2I**). This analysis demonstrated the versatility and reliability of LBL across instruments from different vendors and emphasized the evolving MS technology.

Together, we demonstrated the LBL workflow provides consistent improvement of peptide detection using various multiplex DIA-MS datasets.

### Channel-specific Q-value filtering for quantifying ratios of isotopically labeled peptides

As outlined earlier, LBL performs the elution group scoring across all channels and individually for each channel, leading to the estimation of the group and channel-specific Q-values. These values can be used for quantitative data filtering by e.g., choosing one of the “Group Q-value”, the “Min Q-value”, or the “Max Q-value” options in Spectronaut. In “Min Q-value”, only one of the channels needs to independently pass the Q < 0.01, while in “Max Q-value”, all the channels present in the sample need to pass Q < 0.01 as the most conservative filtering option. We evaluated these three new built-in options using standard 2- and 3-channel SILAC datasets and in the pSILAC experiment of A2780 cells (**Figure 3**).

**Figure 3:**
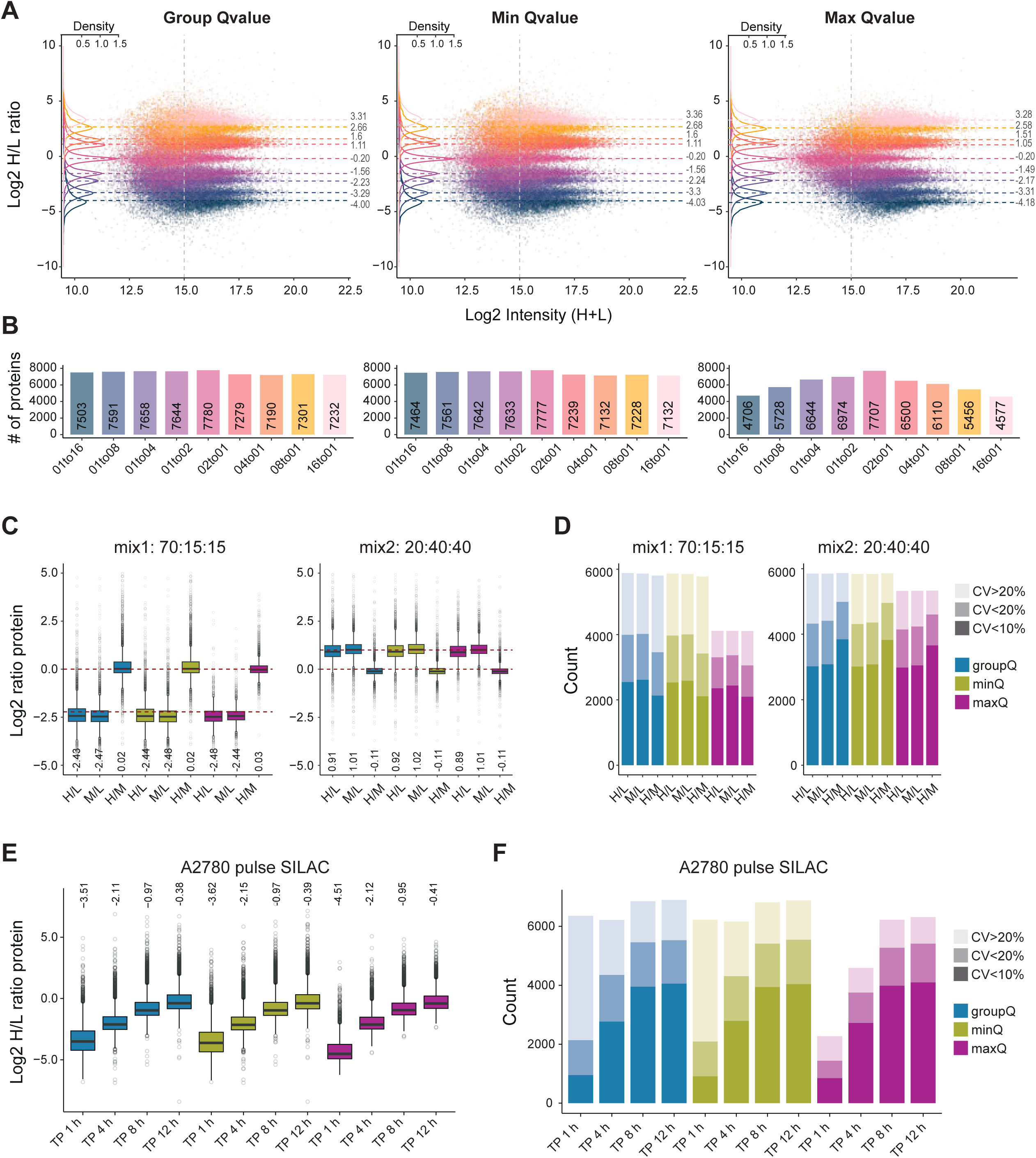
The channel-specific Qvalue filtering for quantifying ratios of isotopically labeled peptides. **(A)** Comparison of protein-level ratio distribution in the A2780 standard dilution samples after different Qvalue quantification filtering for multichannel samples (enabled in SN19); the dashed lines and numbers represent the medians of the data distributions, shown using density plots; the points represent individual values. **(B)** The histograms indicate the number of valid protein-level H/L ratios quantified in the samples depicted in A. **(C)** Comparison of protein-level ratio distribution in the HeLa 3-channel standard sample after different Qvalue quantification filtering. Ratios were calculated between channels as indicated. The dashed lines indicate expected ratios based on sample composition; the numbers represent observed median values. **(D)** Binned protein-level ratio CV based on 3 replicates in the HeLa sample after different Qvalue quantification filtering. **(E)** Comparison of protein-level H/L ratios in the pulse SILAC A2780 samples after different Qvalue quantification filtering. The numbers represent observed median values. **(F)** Binned protein-level ratio CV based on 3 replicates in the A2780 pulse SILAC sample after different Qvalue quantification filtering. groupQ, minQ, and maxQ refer to Group Qvalue, Min Qvalue, and Max Qvalue filtering, which are the quantification settings in the data analysis in SN19.

In the 2-channel dilution series, the “GroupQ” and “MinQ” provided a similar quantification result with comparable numbers of quantified protein-level ratios and accuracy. As expected, the “MaxQ” led to more conservative results filtering with a considerable data loss in the extreme ends of the dilution series, which seemed to be overall intensity dependent (**Figure 3A, 3B**). As expected, MaxQ provided a more stringent filtering that improved quantification precision, as shown by the significantly reduced standard deviations in the H/L ratio distributions, while maintaining similar overall median values. Interestingly, at the precursor level, there is a large overlap between the quantified precursors between the “GroupQ” and “MinQ” filtered results (**Figure S1D**), while GroupQ even consistently identified slightly more precursors (∼3.3% on average), emphasizing the benefit of simultaneously considering all channels.

The analysis of the 3-channel experiment yielded similar conclusions. While the median protein-level ratio values remained relatively consistent across all three filtering options, the “MaxQ” filtering led to an increased precision (**Figure 3C**). The application of “MaxQ” resulted in a significant reduction in the number of quantified proteins (on average 29.9% in mix1 and 9% in mix2), including those with a CV across replicates < 20% (on average 14.1% in mix1 and 5.4% in mix2), indicating the more conservative filtering compromised by the partial loss of high-quality signals.

To benchmark these results against another multiplex DIA-MS workflow, we applied the workflow recommended in plexDIA method with the matrix channel Q-value filtering (Q < 0.01) to analyze both the 2- and 3-channel experiments (**Figure S2**). Notably, the results from plexDIA workflow closely matched those of the “MaxQ” filtering with a higher number of quantified protein-level ratios in “MaxQ” (up to 30% in the 1/16 and 16/1 H/L samples) but slightly better precision using plex-DIA (**Figure S2A**). In the 3-channel sample, we observed a similar trend but a slightly better precision of “MaxQ” (**Figure S2C**). When the analysis was restricted to the same precursors and proteins, the ratio distributions appeared nearly identical (**Figure S2B, S2D**).

Lastly, in the Q-value filtering comparison performed using the real pSILAC A2780 samples, the “MaxQ” filtering significantly reduced the number of quantified protein ratios in the first time point (**Figure 3E, 3F**) and the reported values appeared to have an overall lower median. However, checking the overlapping IDs, the distributions were also largely identical (**Figure S1E**), again suggesting the “MaxQ” reduced noise data points for quantification while discarded sizeable proteins with a good across replicate CV < 20% (32.7% and 13.8% on average in the 1^st^ and 2^nd^-time point; **Figure S3F**).

In summary, in all datasets, both “Group Q-value” and “Min Q-value” options retain higher sensitivity, while the “Max Q-value” is more stringent and delivers improved quantification precision, with the cost of a considerable reduction in quantified proteins.

### *KdeggeR*: A Comprehensive R Package for Proteomic Turnover Analysis

To streamline the analysis of pSILAC-DIA data, herein, we further present ***KdeggeR***, an integrative R package. ***KdeggeR*** offers functions for data import from multiple common raw data processing tools, ensuring compatibility across platforms, followed by data cleaning and quality control steps to prepare the data for analysis (**Figure 4A**). At the precursor level, ***KdeggeR*** allows for the estimation of *k*_loss_ using three different methods, which can then be aggregated to the peptide or protein level. This aggregation is performed by applying a weighted average of precursor-level *k*_loss_ values, with weights determined by the precursor-level fit quality and the number of data points. ***KdeggeR*** also calculates protein degradation rates (*k*_deg_) and half-lives (t_1/2_) using either user-provided or theoretically estimated cell division rates (*k*_cd_), allowing for flexible *k*_deg_ determination. Visualization tools within the package enable users to assess precursor- and protein-level fitting results, as well as conduct comparative turnover analyses between different conditions (see **Methods** for more details).

**Figure 4:**
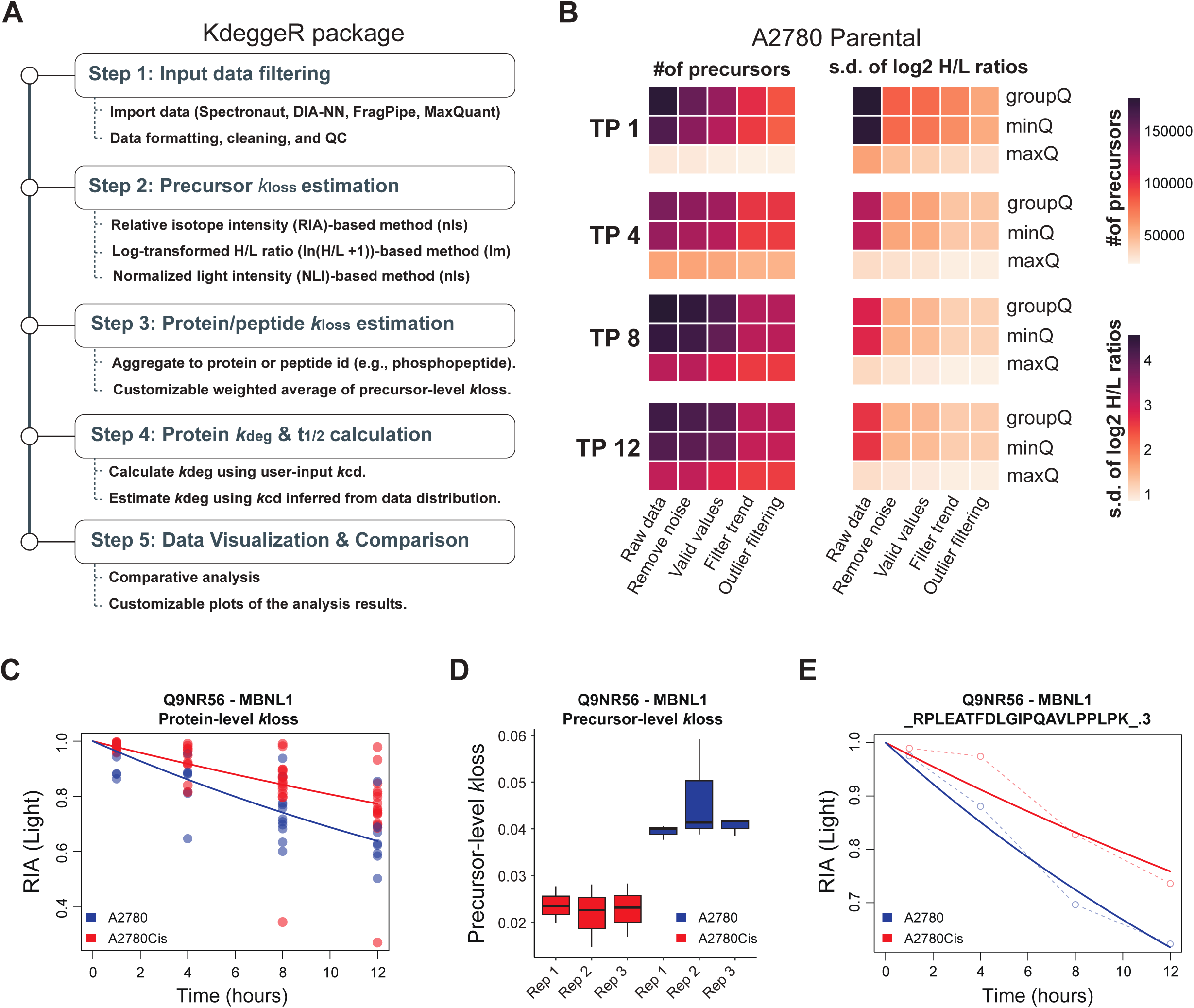
*KdeggeR*, a comprehensive and integrative R package for proteomic turnover analysis. **(A)** The ***KdeggeR*** package streamlines pSILAC data analysis by providing functions for importing data from multiple common raw data processing software tools, data cleaning, and quality control. Next, precursor-level *k*_loss_ can be estimated by three different methods, and protein- and peptide-level *k*_loss_ can be estimated by performing a weighted average of the corresponding precursor-level *k*_loss_ values, considering precursor-level fit quality and/or the number of datapoints. Protein degradation rates (*k*_deg_) and half-life (t_1/2_) are further calculated using cell division rate (*k*_cd_) values provided by the user or by using a theoretical *k*_cd_ value estimated from the *k*_loss_ value distribution. Visualization functions enable inspection of the precursor- and protein-level fitting results and comparative analysis between multiple conditions. **(B)** Demonstration of precursor-level quality filtering in the dynamic SILAC experiment performed in the A2780 cell line. Data were analyzed using the LBL workflow and exported using the Group, Min, and Max Qvalue channel quantification filtering. **(C-E)** An example of MBNL1, a protein with a significantly slower turnover rate in the A2780Cis (resistant) cell line compared to the parental A2780 cell line. **(C)** Protein-level *k*_loss_ fit to all precursor-level data. **(D)** Distribution of precursor-level *k*_loss_ values corresponding to the MBNL1 protein. **(E)** A representative example of precursor-level *k*_loss_ calculation by performing nonlinear least squares (nls) fitting using the relative isotope abundance (RIA) of the light peptide. The plots were visualized using the KdeggeR package.

We utilized the ***KdeggeR*** package to investigate protein turnover regulation between the A2780 and A2780Cis ovarian cancer cell lines ^23^. A triplicate pSILAC experiment was performed with four time points, i.e.,1, 4, 8, and 12 hours (**Figure 1, Lower Right**) and the raw data were processed using LBL. **Figure 4B** demonstrates the precursor-level quality filtering in ***KdeggeR***. We applied a series of filtering criteria considering data completeness and assumptions specific for a pSILAC experiment (see **Methods**).

In addition, outlier values in the early time point (i.e., the first data point) can be removed by performing a linear regression on the log-transformed H/L ratios (ln (H/L +1)) and conducting a statistical test to determine if the first time point significantly deviated from the residual distribution (“Outlier filtering”). It has been established that the first short labeling time point can be critical for precisely determining the turnover rates of short-lived proteins ^9,13^ or modified peptides ^13^. However, due to the lower intensity of heavy-labeled peptides at this initial stage in many pSILAC experiments, the H/L ratios practically tend to exhibit substantially higher noise compared to later time points, which may impact the accuracy of turnover rate quantification. As shown in **Figure 4B**, applying this data filtering approach reduced the standard deviations of H/L ratio distributions, particularly in the first and second time points, while retaining significantly more values with the “GroupQ” filter compared to the more conservative “MaxQ” filter. After fitting precursor-level data with a linear regression of the log-transformed ratios, the majority of curves passed R^2^ > 0.9 (**Figure S3A**). Finally, the precursor and protein-level rates of loss (*k*_loss_) values were estimated using the RIA method and a weighted average (see **Methods**). This resulted in the estimation of 6866 protein *k*_loss_ values on average across the two cell lines and replicates (**Figure S3B**), which were further transformed into *k*_deg_ for the downstream analysis. As an example, ***KdeggeR*** facilitated the identification of MBNL1, a protein exhibiting significantly slower turnover in the A2780Cis (resistant) cell line compared to the parental A2780 line (**Figures 4C-E, S3C**) as visualized by the plotting functions provided within the ***KdeggeR*** package.

Together, ***KdeggeR*** allows for accurate and flexible calculation of protein turnover rates for users without strong bioinformatic background.

### Multi-omics Analysis Reveals Proteomic Buffering via Protein Degradation in a Cisplatin-Resistant Ovarian Cancer Model

Cancer development is often linked to genomic instability and the adaptive evolution of malignant clones. This results in potential genomic alterations conferring selective advantages, including varying responses to chemotherapy. Previous studies, including our own, have demonstrated that cells can exploit the protein degradation system to maintain proteostasis in the face of cell aneuploidy and genomic imbalance ^10,12,24–26^. The A2780 and A2780Cis ovarian cancer cell lines represent a well-established model for studying cisplatin resistance ^23^, with distinct karyotypic abnormalities in both parental and cisplatin-resistant cells documented. These abnormalities have been characterized through genomic ^27,28^ and proteomic ^29^ analyses. However, a comprehensive exploration of protein turnover and its role in driving drug resistance in this drug-resistance model has been lacking.

To investigate the role of protein turnover in regulating genomic imbalance-associated drug resistance, we conducted an integrative multi-omics analysis of the aneuploid A2780 and A2780Cis cell lines. By combining proteomic data—protein abundance and degradation rates (*k*_deg_)—with transcriptomic data from a previous study ^27^, we firstly observed a positive correlation between mRNA and protein log_2_ fold changes (R = 0.661, **Figure S4A**), suggesting a good match between two independent experiments from different laboratories. Furthermore, the correlation between mRNA and *k*_deg_ was weakly positive (R = 0.145, **Figure S4B**), in line with our previous work in HeLa cells ^10,11^, reinforcing the idea that mRNA-*k*_deg_ correlation is a valuable indicator of posttranslational buffering by protein turnover. Next, we used the copy number alteration information (CNA) from a published study ^28^ performed in the same cell lines and mapped the protein-coding genes to the integrated dataset (**Figure 5A**). Reassuringly, the mRNA levels largely followed the expected trend based on the CNA data, and the same trend was apparent in protein abundance data, although the dosage change seemed to be mitigated (**Figure 5A**).

**Figure 5:**
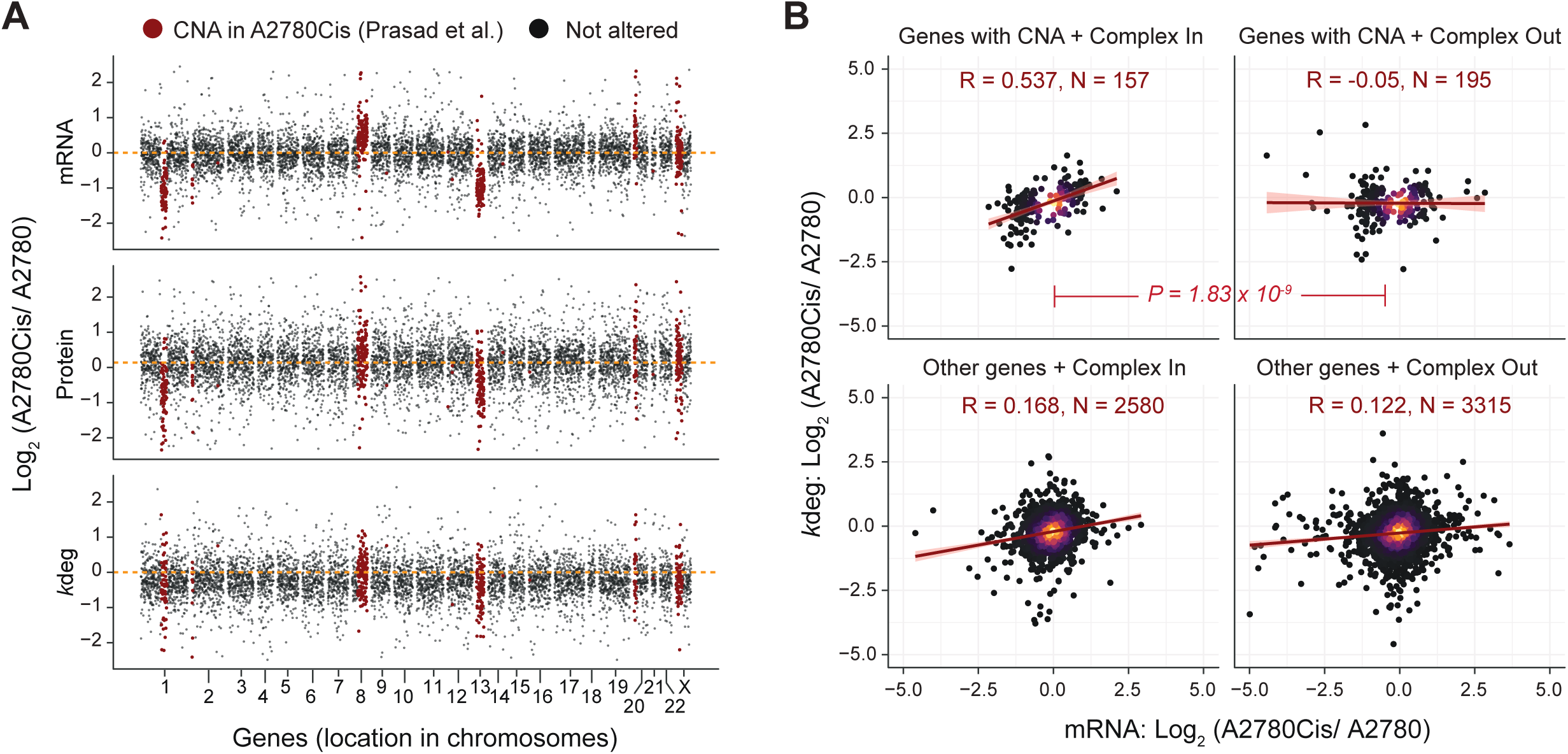
Multi-omic analysis of the cisplatin-resistant model demonstrating proteomic buffering through protein degradation. **(A)** Copy number alterations (CNA) in the A2780 paired cell line model (Parental A2780 vs. A2780Cis) were mapped to transcriptomic data, protein-level abundance data, and protein degradation rate (*k*_deg_) values measured by DIA-MS. The CNA and transcriptomic data were generated by previous studies analyzing the same cell lines (Prasad et al., 2008; Behrman et al., 2021). Relative differences between A2780Cis (resistant) and A2780 (parental) are shown on the y-axis, while the x-axis depicts genes ordered by their chromosome location. Chromosomal regions with CNA are highlighted in red. **(B)** Post-translational buffering through protein turnover of protein complex subunits (Corum 4.0) encoded by genes with reported copy number alterations between the A2780Cis (resistant) and A2780 (parental) cell lines, revealed by mRNA and *k*_deg_ fold change correlations. Statistical analysis was performed using a z-test. Pearson correlation coefficients (R) and the number of proteins (N) are shown. The dark red line represents a linear fit to the data with confidence intervals (pink).

To explore posttranslational buffering in aneuploid cells, we focused on genes affected by CNA and encoding protein complex subunits (Corum 4.0 ^30^). In particular, we plotted the correlation between mRNA and *k*_deg_ which could better inform the proteome buffering existence than protein∼*k*_deg_ correlations as we showed previously ^10^. Remarkably, proteins encoded by CNA-affected genes involved in complexes exhibited a significantly stronger mRNA-*k*_deg_ correlation (R = 0.537, P = 1.83 x 10e^-9^, Fisher’s z-test) than CNA-affected proteins not participating in protein complexes (R = −0.05), providing compelling evidence of large-scale protein complex buffering through protein turnover in this highly aneuploid system. These findings therefore highlight the critical role of protein degradation in buffering against genomic instability, particularly in maintaining the stoichiometry of protein complexes.

### Comprehensive Protein Turnover Analysis Reveals Mechanistic Insights into Cisplatin Resistance in the A2780 Ovarian Cancer Model

In addition to proteome buffering effect, to explore the functional role of protein turnover in the drug-resistant phenotype, we conducted a statistical analysis to identify proteins with significantly altered abundance and degradation rates in the A2780Cis cell line (**Figure S4D, S4E, Table S1**). We identified 1,961 proteins with significant changes in abundance and 1,356 proteins with significantly altered degradation rates, with 407 proteins commonly regulated in both datasets (**Figure 6A**), suggesting both protein abundance and protein turnover regulation are important parts of the drug-resistant phenotype in A2780. Notably, proteins with significantly upregulated *k*_deg_ values overlapped more with the group of significantly downregulated proteins (N = 73, Fisher’s exact test P = 7.11×10e-17) than upregulated (N = 22) (**Figure 6B**), demonstrating a generally coordinated correlation. To corroborate these proteins and their functions, we performed an enrichment analysis using Metascape ^31^, which revealed a densely interconnected cluster of proteins significantly enriched in pathways such as “ATP synthesis coupled electron transport” (P = 7.94e-30) and “Oxidative phosphorylation” (P = 3.16e-31), primarily comprising proteins with increased degradation rates and decreased abundance (**Figure 6D, Table S2**).

**Figure 6:**
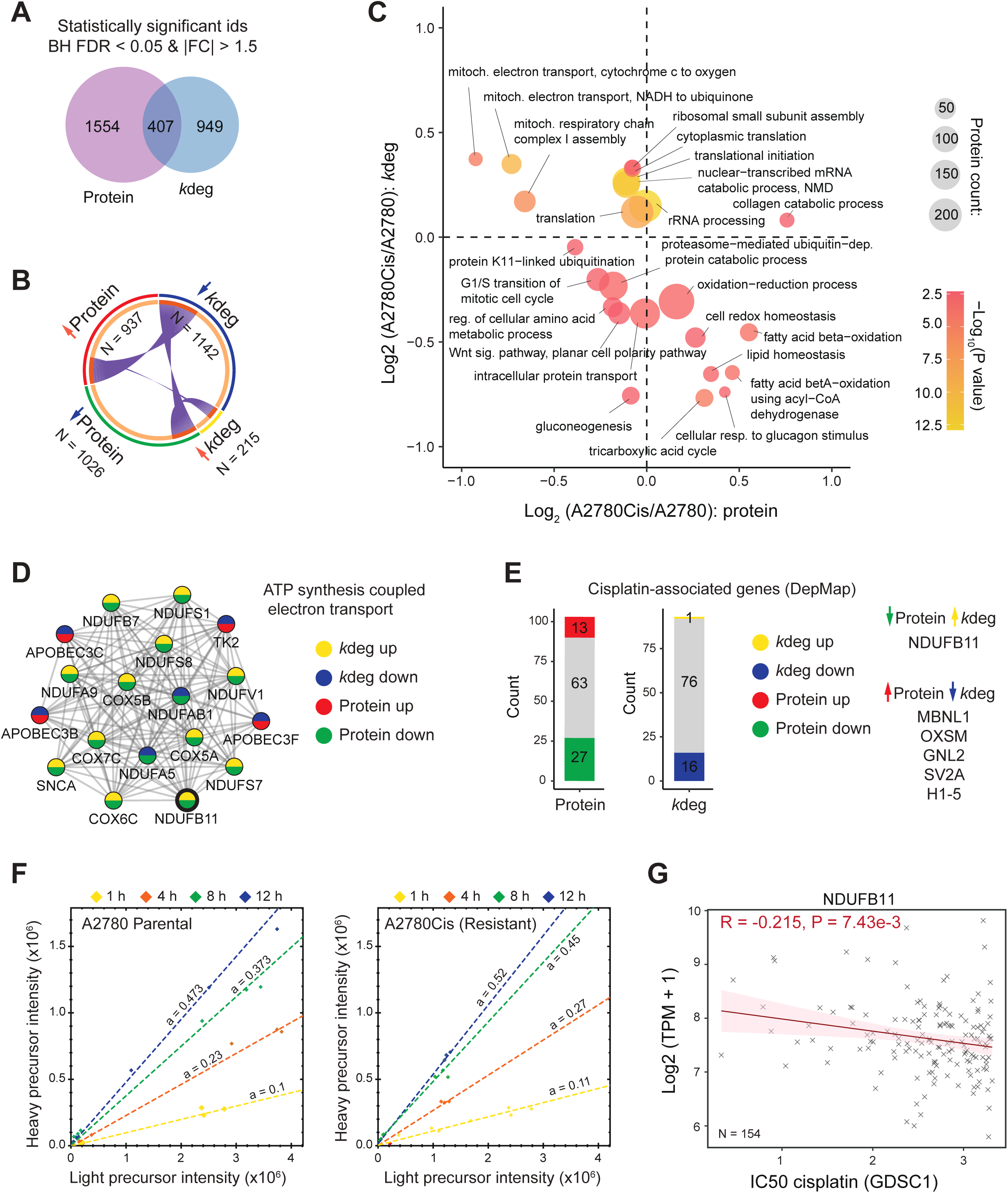
Protein turnover measurement provides biological insights into the drug resistance phenotype in the A2780 cell line model. **(A)** The number of statistically significant protein IDs identified at the protein abundance and degradation levels (Benjamini-Hochberg FDR < 0.05 and absolute fold change of at least 1.5). A moderated t-test was used for statistical analysis. **(B)** The number of significantly up- and down-regulated proteins at the protein abundance and degradation (*k*_deg_) levels, and their overlapping protein identities. The circos plot was generated using Metascape. **(C)** A two-dimensional plot depicts the results of the 2D enrichment analysis of gene ontology biological process (GOBP) functional annotations. Protein abundance and degradation-level relative fold changes between A2780Cis (resistant) and A2780 (parental) cell lines were used for enrichment analysis performed in Perseus. The top 25 GOBP terms based on significance (enrichment P < 0.01 and number of proteins > 9) are shown. The size of the circles represents the number of proteins, while the color represents the −Log10 transformed P value. **(D)** A protein cluster identified by the MCODE algorithm in the protein-protein interaction (PPI) analysis performed using significantly regulated protein IDs (as depicted in B). Node colors represent the respective up- or down-regulation at the protein abundance and degradation levels. **(E)** Genes associated with sensitivity to cisplatin were identified using a custom correlation analysis in the DepMap portal and mapped to the protein abundance and degradation data. The colors represent the respective up- or down-regulation at the protein and protein turnover levels. Proteins with significant regulation at both levels and with opposite trends are shown. **(F)** Precursor-level H/L ratio scatter plots of the NDUFB11 protein in A2780 (left) and A2780Cis (right) cell lines; “a” indicates the slope of the line fitted using a linear fit to the precursor-level data, which can be used as an estimate of the protein-level ratio. **(G)** The negative correlation of NDUFB11 gene expression at the mRNA level with cisplatin IC_50_ based on the GDSC1 dataset indicates that decreased NDUFB11 expression is significantly associated with increased IC_50_ (resistance).

Next, we conducted a two-dimensional gene ontology biological process (GOBP) enrichment analysis, comparing relative changes in both protein abundance and degradation rates between the A2780Cis and A2780 cell lines (**Figure 6C, Table S3**). The correlation between median log_2_ fold changes for significant GOBP terms was overall weakly negative (R = −0.244) as expected. Several processes, such as “TCA cycle”, “lipid homeostasis”, “cell redox homeostasis” or “oxidation-reduction process,” showed increased abundance but reduced turnover, the latter align with mechanisms previously linked to cisplatin resistance ^32,33^. Conversely, proteins involved in “mitochondrial respiratory chain complex I” or “translation” displayed decreased abundance and increased turnover, consistent with findings from the Metascape analysis. Interestingly, terms related to proteasome-mediated protein degradation were downregulated at both proteome and turnover levels.

Additionally, we utilized the DepMap portal ^34^ to identify genes significantly associated with cisplatin sensitivity by examining the correlation between transcript abundance and cisplatin response (N = 414, P < 0.01). Notably, 107 of these genes were mapped to our proteomic datasets, and a substantial number of proteins showed significant regulation, either at the protein abundance or degradation level **(Figure 6E, Table S4**). Proteins that exhibited significant regulation at both levels, but with opposing trends (i.e., coordinated regulation)—upregulation in one accompanied by downregulation in the other— may represent key players involved in mediating drug resistance. Among these, NDUFB11, a mitochondrial electron transport chain protein, was the only one displaying significantly increased degradation **(Figure 6F**) alongside decreased abundance.

A detailed review of DepMap data indeed revealed a negative correlation between NDUFB11 transcript levels and cisplatin sensitivity among a total of 154 cell lines (**Figure 6G**), validating that its reduced abundance, which may be driven by increased turnover, is associated with or contributes to enhanced drug resistance in the A2780Cis cell line. Moreover, MBNL1 and OXSM, which showed increased protein levels and reduced degradation rates in the resistant cells (**Figure S4H**), were positively correlated with cisplatin resistance according to DepMap transcript profiles (**Figures S4F, S4G**), reinforcing their potential roles in mediating the cisplatin resistant phenotype. In conclusion, these findings highlight the strong relationship between protein turnover and cisplatin resistance, with key proteins involved in mitochondrial function, redox homeostasis, and oxidative phosphorylation showing significant regulation.

## Discussion

Multiplex DIA-MS, when integrated with pSILAC ^5–8^, enables precise quantification of protein turnover across multiple time points under various biological conditions ^9–12,14,15,35–37^. This approach is particularly well-suited for time-course experiments, allowing for reproducible profiling of large numbers of peptides. Moreover, recent advances in library-free DIA-MS data analysis driven my machine and deep learning ^19–21^ have dramatically streamlined the process of DIA-MS data identification and quantification by removing the necessity of generating extensive project-specific spectral libraries, making DIA-MS workflows more efficient and scalable for complex proteomic studies.

The LBL in Spectronaut represents an advancement in multiplex DIA-MS data analysis, particularly for experiments involving dynamic isotopic labeling such as pSILAC. Unlike earlier methods that focus on a single channel for signal extraction, the LBL workflow takes a flexible approach by integrating data across all labeling channels. During the peak picking and elution group scoring stages, LBL considers both channel-specific and cross-channel metrics to extract more comprehensive and reliable signals. Moreover, it takes full advantage of the label-free directDIA algorithm in Spectronaut ^20^, effectively eliminating the need for creating a project-specific spectral library. Indeed, we demonstrated that LBL outperformed the ISW in the analysis of multiple labeling datasets, identifying significantly more precursors and protein groups across various conditions and sample compositions. Importantly, we demonstrated improved identification in a standard experiment that included more than two labeling channels, showcasing the potential of LBL to handle different labeling experiments, which theoretically could extend to N channels. This is particularly promising as the interest in multiplexing DIA-MS continues to grow, with new reagents being developed for MS quantification to leverage additional channels in more intricate experimental designs or specialized workflows, such as single-cell proteomics ^17,38^. On the other hand, strategies like ISW or “Spike-in” (with a heavy sample as a reference) may remain more suitable for experiments using spike-in standards ^39^. Finally, we demonstrated that LBL can be effectively utilized across two distinct mass spectrometry platforms—Orbitrap Fusion Lumos and timsTOF Ultra. This adaptability suggests that LBL remains useful even as mass spectrometry technologies evolve, with improvements in speed, resolution, and sensitivity.

Proper FDR control is critical in multiplex-DIA experiments because it ensures the reliability and accuracy of the protein quantifications across multiple labeling channels. LBL leverages machine learning to calculate both cross-channel scores and channel-specific scores. These enable channel-specific FDR filtering of the quantification results three Q-value filtering strategies, “GroupQ”, “MinQ”, and “MaxQ”. Based on our evaluation, the “GroupQ” and “MinQ” (at least one channel Q < 0.01) options offered higher sensitivity by retaining more quantification data, while “MaxQ” (all channels Q < 0.01) provided more stringent filtering, improving precision at the cost of reduced sensitivity. This loss of data in the “MaxQ” setting can be especially impactful in studies focused on low-abundance proteins or studies focusing on peptide-level quantification ^9,13,35^. Researchers may want to consider the choice of filtering strategy based on their specific experimental needs. For studies that prioritize sensitivity, the “GroupQ” or “MinQ” filtering options are more appropriate, while those aiming for the highest level of precision and confidence in their quantification may benefit from using “MaxQ”, albeit with reduced overall data retention. However, it is important to point out that “GroupQ” and “MinQ” results practically cover almost all “MaxQ” results and that Spectronaut offers flexibility with these filtering options, allowing users to tailor their analyses based on their specific experimental requirements. For example, experiments using a booster sample channel might benefit from channel-specific Q-value filtering ^18^.

To provide a comprehensive workflow for the analysis of pSILAC data obtained through multiplex-DIA-MS, we have developed the R package ***KdeggeR***. A few other software tools have been developed previously, among them ***proturn*** ^36^, which offers a user-friendly Shiny app and was primarily designed for pSILAC-TMT experiments; ***JUMPt*** ^40^ calculates protein turnover rates using a differential equation-based model to account for amino acid recycling, and is particularly useful in in vivo studies such as mouse models; ***SPLAT*** enables a more specialized workflow for simultaneous protein localization and turnover analysis (https://lau-lab.github.io/splat/); or ***turnoveR***, which was been designed to work with SVM files and Massacre output (https://github.com/KopfLab/turnoveR). While some of these existing packages can estimate protein degradation rates from pSILAC-DIA data, they sometimes require the user to perform manual data pre-processing. ***KdeggeR*** thus offers an alternative and a more streamlined workflow by handling data import, peptide-to-protein processing, and various data visualization functions within a single package. However, a limitation of the current version of ***KdeggeR*** is its inability to account for amino acid recycling, which can be critical in-vivo systems and is already provided by other software tools such as ***JUMPt*** ^40^.

Finally, we applied the complete workflow to study the potential contribution of protein turnover in the regulation of drug resistance in the A2780 ovarian cancer cell line model ^41^. The cisplatin-resistant cell line A2780Cis was developed by chronic exposure of the parent cisplatin-sensitive cell line A2780 to increasing concentrations of cisplatin, and the development of the drug-resistant phenotype was associated with marked cytogenetic changes ^41^. In this study, we leveraged an integrative multi-omics approach to uncover the mechanisms through which protein degradation helps buffer against genomic imbalances associated with cisplatin resistance. Our focus on genes affected by copy number alterations (CNA) provided compelling evidence of large-scale protein complex buffering ^24^ through protein degradation as observed before ^12^. We observed that proteins encoded by CNA-affected genes involved in complexes exhibited a much stronger mRNA-*k*_deg_ correlation compared to those not involved in complexes, underscoring the importance of turnover dynamics in maintaining the stoichiometry of protein complexes under stress conditions like genomic instability.

In terms of cisplatin resistance, our analysis of the A2780Cis cell line revealed significant changes in both protein abundance and degradation rates. Proteins with upregulated *k*_deg_ values tended to have reduced abundance, indicating that specific degradation mechanisms potentially critical for maintaining drug resistance. Notably, key pathways such as oxidative phosphorylation and ATP synthesis were significantly enriched among proteins with increased degradation rates and reduced abundance, suggesting that mitochondrial function might be an important factor in the cisplatin resistance phenotype. Moreover, processes such as cell redox homeostasis, and the TCA cycle were associated with increased abundance and decreased turnover. Conversely, pathways related to mitochondrial respiratory chain complex I and translation showed decreased abundance and increased turnover, highlighting a potential disruption in energy metabolism and protein synthesis machinery in the resistant cells. Several studies have highlighted the potential role of alterations in drug uptake, enhanced DNA repair pathways such as nucleotide excision repair (NER), and cytosolic drug inactivation as the main contributing factors to cisplatin resistance ^23,42^. Other studies have suggested that drug inactivation in the cytosol by the cell redox system involving metallothionein ^32^ and/or glutathione ^33^ may contribute to cisplatin resistance. This aligns with our observation of an enhanced cellular redox system in the cisplatin-resistant A2780Cis cell line.

Finally, we used the DepMap portal ^34^ to identify genes associated with cisplatin sensitivity, revealing several proteins with significant regulation at both the protein abundance and degradation levels. Of particular interest was **NDUFB11**, a mitochondrial electron transport chain protein, which exhibited increased degradation and decreased abundance. This suggests that altered turnover of mitochondrial proteins and alterations in mitochondrial functions may contribute to drug resistance. Similarly, **MBNL1** and **OXSM**, which showed increased protein levels and reduced degradation rates, may represent important contributors to the cisplatin-resistant phenotype. The observed synergetic mechanisms, such as increased protein degradation paired with reduced abundance, underscore the critical role of turnover dynamics in maintaining cellular proteostasis under aneuploidy and drug stress.

In conclusion, our integration of multiplex DIA-MS with pSILAC and the development of the LBL workflow in Spectronaut significantly enhance protein turnover quantification across multiple channels. These advancements streamline complex proteomic studies and provide valuable insights into mechanisms such as cisplatin resistance.

## Acknowledgement

Y.L. thanks the support from the National Institute of General Medical Sciences (NIGMS), National Institutes of Health (NIH) through Grant R01GM137031 to Y.L. We would like to thank Diego Assis and Matthew Willets from Bruker Life Sciences, Billerica, MA for performing the LC-MS analysis using the timsTOF Ultra platform.

## Author Contributions

B.S. analyzed the MS data, performed the bioinformatics analysis, and prepared most illustrations for the figures. W.L. prepared all the samples and performed MS measurements. O.B., T.G, and L.R. contributed to the software development facilitating multiplex DIA-MS data analysis. P.L.G. and B.S. developed the KdeggeR package. Y.L. secured funding and supervised the study. B.S. wrote the first version of manuscript. All authors contributed to the writing of the manuscript.

## Methods

### Key resources table

| REAGENT or RESOURCE | SOURCE | IDENTIFIER |
| --- | --- | --- |
| **Biological Samples** |  |  |
| A2780 | Sigma | Cat #: 93112519 |
| A2780Cis | Sigma | Cat #: 93112517 |
| Fibroblast from Skin (#54) | Coriell Institute |  |
| Fibroblast from Skin (#55) | Coriell Institute |  |
| **Chemicals and Reagents** |  |  |
| MEM, NEAA, no glutamine | Gibco (Thermo Fisher Scientific) | Cat #: 10370088 |
| MEM for SILAC | Thermo Scientific | Cat #: PI88368 |
| RPMI 1640 medium | Gibco (Thermo Fisher Scientific) | Cat #: 11875093 |
| RPMI 1640 Medium for SILAC | Gibco (Thermo Fisher Scientific) | Cat #: 88365 |
| Fetal bovine serum | Gibco (Thermo Fisher Scientific) | Cat #: A5256701 |
| Fetal bovine serum (dialyzed) | Gibco (Thermo Fisher Scientific) | Cat #: 26400044 |
| GlutaMAX™ Supplement | Gibco (Thermo Fisher Scientific) | Cat #: 35050061 |
| HyClone Penicillin-Streptomycin | Cytiva | Cat #: SV30010 |
| PBS, pH 7.4 | Gibco (Thermo Fisher Scientific) | Cat #: 10010023 |
| Trypsin-EDTA (0.25%), phenol red | Gibco (Thermo Fisher Scientific) | Cat #: 25200056 |
| Heavy L-Arginine-HCl (13C6, 15N4, purity >98%) | Cortecnet | Cat #: CCN250P1 |
| Heavy Lysine-2HCl (13C6, 15N2, purity >98%) | Cortecnet | Cat #: CCN1800P1 |
| L-Proline | Sigma | Cat #: P0380 |
| Urea | Sigma | Cat #: U5378 |
| Halt™ phosphatase inhibitor | Thermo Scientific | Cat #: 78428 |
| cOmplete™ protease inhibitor cocktail | Roche | Cat #: 11697498001 |
| Ammonium bicarbonate | Sigma | Cat #: A6141 |
| Bio-Rad protein assay kit | Bio-Rad | Cat #: 5000002 |
| Sequencing-grade modified trypsin | Promega | Cat #: V5113 |
| Formic acid | Thermo Scientific | Cat #: 85178 |
| Trifluoroacetic acid, LC-MS grade | Thermo Scientific | Cat #: 85183 |
| DTT | Sigma | Cat #: D0632 |
| IAA | Sigma | Cat #: I1149 |
| 96-Well Macro SpinColumn | Harvard Apparatus | Cat #: 74-5657 |
| 96-Well Spin Column | The Nest Group, Inc | Cat #: 8003476378 |
| ReproSil-Pur,120A, C18-AQ,1.9 um resin | Dr. Maisch | Cat #: r119.aq |
| PicoFrit LC-MS column | New Objective | Cat #: PF360-75-10-N-5 |
| Cisplatin | Sigma | Cat #: C2210000 |
| Cell Lifter, Fixed 2.0cm Blade | RPI | Cat #: 162424 |
| 2 ml eppendorf tube | Eppendorf | Cat #: 022363433 |
| Corning® 100 mm TC-treated Culture Dish | Corning | Cat #: 430167 |
| Cytoone 6-well Plate with Lid, TC-treated | USA Scientific | Cat #: CC7682-7506 |
| Acetone | Fisher Scientific | Cat #: A18P4 |
| LC-MS grade Acetonitrile | Thermo Scientific | Cat #: 85188 |
| Water, Optima™ LC/MS Grade, Fisher Chemical™ | Fisher Scientific | Cat #: W64 |
| Methanol | Fisher Scientific | Cat #: A412P4 |
| 0.1% Formic Acid (v/v) in Water, LC-MS Grade | Fisher Scientific | Cat #: PI85171 |
| Acetonitrile with 0.1% Formic Acid (v/v), Optima™ LC/MS Grade | Fisher Scientific | Cat #: LS120500 |
| Ethyl alcohol, Pure | Sigma | Cat #: E7023 |
| **Equipment** |  |  |
| VialTweeter sonicator | Hielscher-Ultrasound Technology | Cat #: UP 2000St |
| SpeedVac | Thermo Scientific |  |
| Nanodrop | Thermo Scientific |  |
| Microplate Reader | BioTek |  |
| ThermoMixer | Thermo Scientific | Cat #: 5382000023 |
| **Instruments** |  |  |
| Column heater controller | Sonation GmbH, Biberach | Cat #: PRSO-V1 |
| Easy nLC 1200 system | Thermo Scientific | Cat #: 1200 |
| Orbitrap Tribrid Lumos Mass Spectrometer | Thermo Scientific |  |
| **Deposited Data** |  |  |
| A2780 standard dilution sample set | PXD021922 |  |
| Fibroblast pSILAC sample set | x |  |
| A2780 cells, parental and cisplatin-resistant, pSILAC and total proteome | x |  |
| **Software and Algorithms** |  |  |
| Spectronaut™ Professional+ | v19 | Biognosys AG |
| DIA-NN | v1.9 | Demichev et al (2020), Derks et al (2023) |
| Perseus | v1.6.14.0 | Tyanova et al (2016) |
| Rstudio | v4.3.1 | R core team (2024) |

### Sample sets

#### A2780 standard dilution sample set (2-channel SILAC)

This sample set was measured in our previous study ^15^. Ovarian cancer cell line A2780 was cultured for at least eight passages in media containing^13^C_6_^15^N_4_-Arg and ^13^C_6_^15^N_2_-Lys to reach >99% labeling efficiency (as evaluated by MS) ^15^. The sample set included the following H/L dilutions: 1:16, 1:8, 1:4, 1:2, 1:1, 2:1, 4:1, 8:1, and 16:1. The detailed sample preparation and LC-DIA-MS protocol can be accessed in the original protocol and the raw files provided at ProteomeXchange (**PXD021922**). In brief, a 4-hour method consisted of an MS1 survey and 33 MS2 scans of variable windows ^9^ on an Orbitrap Fusion Lumos Tribrid mass spectrometer (Thermo Scientific). ***HeLa standard dilution sample set (3-channel SILAC).*** This sample set included data from a published study ^22^ and was downloaded from ProteomeXchange (**PXD039578**). The cells were grown in high glucose DMEM, dialyzed fetal bovine serum (Gibco), and heavy-(^13^C_6_^15^N_4_-Arg, ^13^C_6_^15^N_2_-Lys), intermediate (^13^C_6_-Arg, D_4_-Lys) or light isotope-containing Lysine, Arginine for 10 days. The H:M:L composition of the mix 1 sample was made to be 15:15:70, and the composition of the mix 2 sample was 40:40:20. These samples were analyzed in triplicates using the LC-DIA-MS as described in the original paper (MS2-optimized; see ^22^ for details). In brief, a 105-min method consisted of an MS1 survey and 26 MS2 scans of equally sized windows of 23.3 m/z on a QExactive HF mass spectrometer (Thermo Fischer Scientific).

#### Fibroblast pSILAC sample set (2-channel pulse SILAC)

The skin fibroblast cell lines were purchased from the Coriell Institute for Medical Research. The two cell lines are referred as cell line #54 and cell line #55 in the current manuscript, respectively. The cells were cultured at 37 ℃, and humidified 5% CO_2_ in complete MEM medium supplemented with L-glutamine, 15% fetal bovine serum, and penicillin-streptomycin. Cells were seeded on 6-well dishes in a complete growth medium at a density of 15,000 cells per cm^2^, and after 24 hours, the cells were washed and subjected to pulse SILAC labeling for 1, 4, 8, 12, and 24 hours. The SILAC MEM medium was supplemented with 15% of dialyzed FBS, penicillin-streptomycin, L-proline (200 mg/L), and ^13^C_6_^15^N_4_-Arg and ^13^C_6_^15^N_2_-Lys. The dishes were washed with PBS, snap-frozen in liquid nitrogen, and the cells were scraped into 100 µL of cell lysis buffer containing 10 M urea/ 100 mM ammonium bicarbonate, cOmplete™ protease inhibitor cocktail and the Halt phosphatase inhibitors. The collected samples were snap-frozen and stored at −80 °C before further processing.

#### A2780 cells, parental and cisplatin-resistant (2-channel pulse SILAC)

The A2780 (parental cell line) and A2780Cis (cisplatin-resistant cell line) were cultured in the RPMI media supplemented with 2 mM glutamine, 10% FBS, and penicillin-streptomycin. Additionally, the A2780Cis cell line was cultured in the presence of 1 µM cisplatin. After switching to SILAC heavy medium (^13^C_6_^15^N_4_-Arg and ^13^C_6_^15^N_2_-Lys), both cell lines were harvested in a triplicate experiment at 1, 4, 8, and 12 hours of labeling. Additionally, a triplicate sample was harvested at time point 0 to analyze the total proteomes. The dishes were washed with PBS, snap-frozen in liquid nitrogen, and the cells were scraped into 200 µL of cell lysis buffer containing 10 M urea/ 100 mM ammonium bicarbonate, cOmplete™ protease inhibitor cocktail and the Halt phosphatase inhibitors. The collected samples were snap-frozen and stored at −80 °C before further processing.

### Protein extraction and digestion

Cell pellets in lysis buffer were thawed and sonicated at 4°C for 1 minute twice using a VialTweeter device (Hielscher-Ultrasound Technology) ^11^. Afterward, the samples were centrifuged at 20,000 × g for 1 hour to separate insoluble materials. Protein concentrations in the resulting supernatant were measured using the Bio-Rad protein assay. Each protein sample was diluted to a final concentration of 2 μg/μl, reduced with 10 mM DTT at 56°C for 1 hour, and alkylated with 20 mM IAA in the dark at room temperature for 1 hour. Reduced and alkylated proteins underwent precipitation-based digestion ^43^ or in-solution digestion. For the precipitation-based digestions (all A2780 samples), five volumes of a cold precipitation solution (50% acetone, 50% ethanol, and 0.1% acetic acid) were added to the protein mixture, and the samples were stored at −20°C overnight. The precipitated proteins were collected by centrifugation at 20,000 × g for 40 minutes, washed with cold 100% acetone, and centrifuged again under the same conditions. Following acetone removal, residual acetone was evaporated in a SpeedVac. The proteins were then digested overnight at 37°C with sequencing-grade porcine trypsin at a 1:20 enzyme-to-substrate ratio in 300 μl of 100 mM ammonium bicarbonate. For the in-solution digestion (fibroblast samples), the samples were diluted five times with 100 mM ammonium bicarbonate prior to the addition of trypsin in 1:20 enzyme-to-substrate ratio. The peptide mixture was acidified with formic acid and desalted using C18 columns (MarocoSpin Columns, NEST Group INC.) according to the manufacturer’s instructions. The final peptide yield was quantified using a Nanodrop (Thermo Scientific).

### Mass Spectrometry Measurements

#### Orbitrap Fusion Lumos platform

For LC-MS analysis, 1.5 μg of the peptide mixture was analyzed as previously described ^11,44^. Peptide separation was carried out using an EASY-nLC 1200 system (Thermo Scientific) with a self-packed PicoFrit column (New Objective, Woburn, MA, USA; 75 μm × 50 cm) containing ReproSil-Pur 120A C18-Q 1.9 μm resin (Dr. Maisch GmbH, Ammerbuch, Germany). Peptides were eluted over a 150-minute gradient using buffer B (80% acetonitrile, 0.1% formic acid) from 5% to 37%, with buffer A (0.1% formic acid in water) as the corresponding solvent. The flow rate was set to 300 nl/min, and the column was maintained at 60°C using a column oven (PRSO-V1; Sonation GmbH, Biberach, Germany). The separated peptides were analyzed on an Orbitrap Fusion Lumos Tribrid mass spectrometer (Thermo Scientific) equipped with a NanoFlex ion source, with a spray voltage of 2000 V and a capillary temperature of 275°C. The DIA-MS method included an MS1 survey scan followed by 33 MS2 scans with variable windows, as described previously ^20,45^. The MS1 scan range was 350–1650 m/z with a resolution of 120,000 at m/z 200. The MS1 AGC target was set to 2.0E6, with a maximum injection time of 100 ms. For MS2, the resolution was set to 30,000 at m/z 200, with a normalized HCD collision energy of 28%. The MS2 AGC target was 1.5E6, and the maximum injection time was 50 ms. The default peptide charge state was set to 2. Both MS1 and MS2 spectra were recorded in profile mode.

#### timsTOF Ultra platform (fibroblast pSILAC dataset)

Peptides (130 ng) were separated within 52-minute ACN gradients on a 25cm x 75µm column (Ion Opticks) using a nanoElute2 LC. The LC system was connected via a CaptiveSpray Ultra source to trapped ion mobility – quadrupole time-of-flight MS (timsTOF Ultra, Bruker Daltonik). The MS was operated in dia-PASEF mode ^46^ with 3 PASEF mobility scans, each with 20 DIA variable windows (a “20 × 3” method; Bruker Daltonics) ^47^.

### Raw data processing

#### Label-free DIA-MS data analysis (A2780 and A2780Cis)

The label-free data analysis was performed in Spectronaut v19 using directDIA+ against a human SwissProt sequence database (N = 20,399 entries, downloaded in September 2022) using the default settings ^20^. Briefly, the Trypsin/P was used as a cleavage rule with up to 2 missed cleavages; “Carbamidomethyl(C)” was set as a fixed modification, and “Acetyl (Protein N-term)” and “Oxidation(M)” were set as variable modifications; Top3-6 Best N Fragments per peptide were enabled. The precursor Q-value and the experiment-wide protein Q-value were set to 0.01, and the run-wise protein Q-value was set to 0.05. The quantification was performed on the MS2 level, and the cross-run normalization was enabled. The peptide and protein quantification were performed using max Top 3 precursors and Top 3 stripped peptide sequences, respectively. The “Minimum Log2 Precursor Quantity” was set to 3.

#### Multiplex DIA data analysis using the “Labeled workflow” (LBL)

The multiplex DIA data analysis was performed in Spectronaut v19 using the library-free “Labeled” workflow. The analysis was performed using directDIA+ against a human SwissProt sequence database (N = 20,399 entries, downloaded in September 2022) using the default settings with modifications as described below. The search parameters were the same in all datasets across MS platforms.

In the Pulsar Search: the Trypsin/P was used as a cleavage rule with up to 2 missed cleavages; the labeling was set to two channels with no labels specified in Channel 1 and “Arg10” and “Lys8” specified in Channel 2; “Carbamidomethyl(C)” was set as a fixed modification, and “Acetyl(Protein N-term)” and “Oxidation(M)” were set as variable modifications; in the Workflow tab, the “In-Silico Generate Missing Channels” option was enabled with “label” as a Workflow; in the Result Filters tab, Top3-6 Best N Fragments per Peptide were used, and the “Overlapping between Channels” was enabled to exclude fragments shared between channels for the accurate estimation of channel-specific FDR.

In the DIA Analysis: in the Identification tab, the precursor Q-value and the experiment-wide protein Q-value were set to 0.01, the run-wise protein Q-value was set to 0.05; in the Quantification tab, the Multi-Channel Q-value filter was either set to “*Group Q-value*”, “*Max Q-value*”, or “*Min Q-value*” to evaluate channel-specific Q-value filtering options. The quantification was performed on MS2 level, and the cross-run normalization was enabled. The “Exclude All Multi-Channel Interferences” option was enabled. The peptide and protein quantification were performed using max Top 3 precursors and Top 3 stripped peptide sequences, respectively. The “Minimum Log2 Precursor Quantity” was set to 3. In the Workflow tab, the “Multi-Channel Workflow Definition” was set to “Labeled”.

#### Multiplex DIA data analysis using the “Inverted Spike-In” workflow (ISW)

For the “Inverted Spike-In analysis, the “Multi-Channel Workflow Definition” was set to “Spike-In” and both “Inverted” and “Reference-based Identification” were enabled in Spectronaut v19. Other parameters were kept as described in the section above.

#### Multi-Channel Experiment Processing and Scoring in Spectronaut

Spectronaut organizes all channels corresponding to a given peptide into “ElutionGroups,” representing a group of peptide precursors expected to elute simultaneously. Extracted ion chromatograms (XICs) are obtained for each group from the relevant MS2 scans. The peak-picking strategy depends on the specific multi-channel processing mode selected. By default, Spectronaut utilizes the “Labeled” workflow for multi-channel ElutionGroups, in which XIC peak picking is performed across all channels in a combined fashion. Each peak is assigned scores based on both channel-specific and cross-channel features, for both MS1 and MS2 data.

The final score per ElutionGroup is determined during the machine learning step, with the strategy being workflow dependent. In the “Labelled” (LBL) workflow, all channels, along with their specific and cross-channel scores, are considered collectively in the machine learning process, allowing for dynamic adaptation to systematic changes in channel ratios which are common in pSILAC experiments. These scores are then used to compute the “Group Q-value” for each ElutionGroup. Additionally, channels are scored independently based on channel-specific metrics, which are used to determine channel-specific Q-values.

Note, since this filtering is performed at the precursor level, the quantification of an elution group (and thus the ratios between the channels) will be identical for those precursors passing the FDR filtering by multiple options.

### Spectronaut results export and report processing

#### Label-free DIA-MS data (A2780 and A2780Cis)

For the total proteome analysis, the data were exported using the protein pivot report using PG.ProteinGroups as a unique protein id and the PG.Quantity as a quantification column.

#### Multiplex DIA-MS data

The precursor/elution group (EG) level pivot report was exported from Spectronaut v19. The “EG.Channel1”and “EG.Channel2” (for a 2-channel experiment), and “EG.Channel3” (for a 3-channel experiment) quantities were used as quantification values, and the “EG.PrecursorId” column as the unique precursor id column. For protein-level quantification of the multiplex DIA-MS data, the precursor-level ratios were estimated first as the ratios between the channels, and then the protein-level ratios were calculated as a median value of all precursor-level ratios corresponding to a unique protein id (“PG.ProteinGroups”). In all replicate experiments (HeLa 3-channel, A2780 pSILAC), the replicates were aggregated to obtain an average ratio by calculating a mean and CV of non-transformed ratio values after filtering for precursor values quantified in all 3 replicates.

#### Multiplex DIA-MS data analysis in DIA-NN

To benchmark the multiplex DIA-MS analysis in Spectronaut, we also analyzed the data using DIA-NN (version 1.9) ^19^ using the plexDIA recommended workflow ^17^ and largely followed the parameters used previously to analyze the 3-channel HeLa standard sample experiment ^22^. A predicted spectral library was generated using the default settings from the same FASTA file used for Spectronaut searches and the same fixed and variable modifications. For the raw data analysis, the default settings were used, along with additional commands necessary to analyze a plexDIA experiment (https://github.com/vdemichev/DiaNN). Specifically, the SILAC channels were registered, depending on the 2- or 3-channel experiment, corresponding to Lysine and Arginine mass shifts: Lys (+4.025107 Da), Lys8 (+8.014199 Da), Arg6 (+6.020129 Da), Arg10 (+10.008269 Da). Retention time translation between peptides within the same elution group was enabled. Both the first 13C-isotopic and monoisotopic peaks were included for quantification, with MS1 deconvolution level set to 2. Peptide lengths ranged from 7 to 30 amino acids, precursor charge states ranged from 1 to 4, and the precursor mass-to-charge (m/z) range was set between 300 and 1800, with fragment ion m/z range from 200 to 1800. The precursor false discovery rate (FDR) was set to 1%. Precursor matrix output tables were filtered for FDR < 0.01, as well as for channel-specific (--matrix-ch-qvalue) and translated q-values < 0.01 (--matrix-tr-qvalue). The match-between-runs (MBR) function in DIA-NN was enabled. The precursor-level matrices were used for the downstream analyses (“report.pr_matrix_channels.tsv”).

### Determination of protein degradation rates from the pulse SILAC experiments

Protein degradation rates reported in this manuscript were calculated using the *KdeggeR* package following an algorithm based on the nls fitting in the relative light isotope abundance values (RIA_Light_) at the precursor level as described in detail previously ^5,10,11,48–50^ and subsequent averaging to the protein-level rates of loss and degradation rates. The main steps are described below together with the description of additional options and functionalities of the package. The package will be provided via github.

#### Data import, formatting, and filtering

The precursor-level report from Spectronaut was imported and the channel intensity values were filtered to remove low-intensity signals (e.g., at log_2_-transformed intensity < 8). Note, that this filtering significantly improves data quality in our datasets and is recommended to perform in the multi-channel data analysis in Spectronaut by default, using the “Minimum Log2 Precursor Quantity” quantification filter. Next, the H/L ratios were calculated and further filtered based on i) valid values (e.g., at least 2 in time points 4, 8, and 12 for the A2780 datasets), ii) increasing trend over the time points, and iii) outliers were detected in the first time point based on the identification of significant outliers using linear regression. To do so, we fit a linear model using log-transformed H/L ratios (ln(H/L +1)) from time points 4, 8, and 12. We then calculated residuals of the fit per each time point, including the first time point. Grubb’s test was used as a statistical test to detect significant outliers from the residual distribution in time point 1.

#### Estimation of precursor-level k_loss_ values using the RIA method

At each time point, the amount of heavy (H) and light (L) precursor was extracted and used to calculate the relative isotopic abundance RIA_t_.

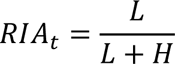

The value of RIAt changes over time as unlabeled proteins are gradually replaced by heavy-labeled proteins throughout the experiment. This occurs because of cell division, which dilutes the unlabeled proteins, and the natural turnover of intracellular proteins, where the loss rate can be described by an exponential decay process.

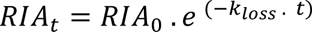

Where RIA_0_ denotes the initial isotopic ratio and *k*_loss_ the rate of loss of unlabeled protein. We assumed RIA_0_ = 1, as no heavy isotope was present at t = 0, thus the value of RIA_t_ will decay exponentially from 1 to 0 after infinite time and used nonlinear least-squares estimation to perform the fit. As discussed before ^5^, these assumptions may reduce measurement error, especially at the beginning of the experiment, where isotopic ratios are less accurate.

#### Estimation of precursor-level k_loss_ values using the NLI method

A simpler approach to determine *de facto* protein degradation rates is to directly calculate the rate of loss from the light peptide intensities. The light peptide intensities need to be normalized using median channel sums to calculate the normalized intensity values (NLI). Then, the light precursor rate of loss can be modeled using the same model and assumptions as in the case of the RIA-based modeling. As we reported previously, the NLI and RIA method results are strongly correlated, however, the NLI method tends to have higher variability ^10^.

#### Estimation of precursor-level k_loss_ values using the HOL method

The heavy proteins are synthesized over time, leading to an increasing H/L ratio. This process is exponential because the heavy proteins are gradually replacing the unlabeled (light). The H/L ratios are linearized by log-transformation and the rate of incorporation of the heavy label is then estimated from a linear model.

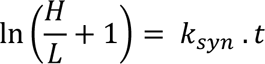

In the steady-state condition, the rates of protein synthesis and degradation reach equilibrium. This means that the rate at which new heavy-labeled proteins are synthesized must be balanced by the rate at which proteins are degraded or turned over (*k*_loss_).

#### Estimation of protein-level k_loss_ values

Protein-level *k*_loss_ values can be calculated by performing a weighted average of the selected fit (e.g., RIA only) or their combination (e.g., RIA and NLI). The number of data points used to estimate precursor-level *k*_loss_, the variance of the fit, or both can be used as weights.

#### Calculation of protein-level k_deg_ values

Protein degradation rates are estimated by subtracting the cell division rates (*k*_cd_) to correct for the protein pool dilution caused by the exponential cell division.

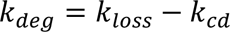

However, based on our experience, the cell division rates tend to be very variable between different experiments and thus the precision and accuracy tend to be low. Therefore, we decided to use a *k*_cd_ derived from the distribution of the *k*_loss_ values by assuming that most *k*_deg_ values should be positive after the correction. We herein suggest a value (*k*_perc_) by subtraction of which only 1% of *k*_deg_ values would be negative that the users may be able to estimate the *k*_deg_ values in cell culture derived datasets.

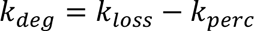

Optionally, protein half-lives from the degradation rate constant using the following formula. ln (2)

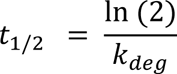

Note, for the results presented in this manuscript, we used the precursor *k*_loss_ estimation using the RIA method, and then calculated the protein *k*_loss_ as a weighted average of the precursor-level data using both number of data points and variance as the weights of the fit. The *k*_deg_ values were calculated by subtracting the theoretical *k*_perc_ from the *k*_loss_ values.

### Downstream bioinformatic analysis of the drug-resistance model experiment

#### aCGH dataset

Array comparative genomic hybridization (aCGH) data were downloaded from a from a previous study (their Supplementary Table 1) ^28^. Protein-coding genes mapping to the regions with gene copy number alterations (CNA) between the A2780 (parental) and A2780Cis (cisplatin resistant) provided in the table were identified by mapping those regions to the homo sapiens Ensembl genome using the bioMart R package.

#### RNA-seq dataset

The RNA sequencing results were downloaded from the GEO under accession **GSE173201** which were published in a previous study ^27^. A table containing TPM normalized counts (GSE173201_norm_counts_TPM_GRCh38.p13_NCBI.tsv) was used for all analyses presented in this study. The data were filtered for transcript matching to protein-coding genes and filtered for genes with at least 2 valid non-zero values (out of 3 replicates) in each condition. The TPM data were transformed using the limma::voom() function before further statistical analysis.

#### Proteome abundance analysis

The protein-level pivot report tables were exported from Spectronaut v19, log_2_-transformed, and normalized using the limma::normalize.cyclic.loess() function ^51^. The data were filtered to only contain proteins quantified in at least 2 out of three experimental replicates before the statistical analysis.

#### Protein degradation analysis

The *k*_deg_ values exported from the KdeggeR package were log_2_-transformed and filtered to only contain proteins quantified in at least 2 out of three experimental replicates before the statistical analysis.

#### Statistical analysis

The statistical analysis was performed using the limma ^52^ R package following the standard pipeline of lmFit(), contrasts.fit(), and eBayes(). The limma results were corrected for multiple testing using FDR correction using the Benjamini-Hochberg method. Cutoffs of FDR < 0.05 and an absolute fold change > 1.5 were used to report significantly regulated features in the RNA-seq, protein abundance, and protein degradation datasets. The moderated log2-transformed fold changes exported from the results were used in all downstream analyses.

#### Datasets integration and correlation analysis

The datasets were integrated based on unique gene symbols. The absolute correlation analysis was performed for the A2780Cis cell line using ids successfully quantified in all 3 layers (N = 6,221) and using average log_2_ TPM, log_2_ protein intensity, and log_2_ *k*_deg_; and Spearman’s Rho was reported. The relative correlation analysis was performed using moderated log_2_ fold changes using ids with valid log_2_ fold change in all 3 layers (n = 6,203); and Pearson’s correlation was reported. For the *k*_deg_∼mRNA analysis presented in Figure 5, the data were split into four groups based on two parameters to perform a correlation analysis and statistical analysis using a Fisher’s z-test. As for the first parameter, genes affected by CNA were identified all protein-coding genes identified based on the aCGH data, excluding genes encoded by the X chromosome. As for the second parameter, the genes were further split based on the participation of the encoded protein in protein complexes as retrieved from the Corum 4.0 database ^30^.

#### Functional Enrichment analyses

A multiple gene list enrichment analysis was performed using the Metascape web interface (https://metascape.org) ^31^ to perform the functional enrichment analysis. Four lists of protein IDs were provided based on the results of the above statistical analysis (protein “up”, protein “down”, *k*_deg_ “up”, and *k*_deg_ “down”), and the default parameters were used. All protein-protein interactions (PPI) from the STRING database ^53^ between the four lists of proteins were used to generate a PPI network followed by the Molecular complex detection (MCODE) algorithm ^54^ to identify densely inter-connected clusters in the PPI network following a gene ontology (GO) enrichment analysis. The resulting color-coded protein-protein interaction networks were further processed in Cytoscape (v 3.10.1) to generate figures presented in Figure 6. Metascape output was also used to generate the Circos ^55^ plot in Figure 6. The 2D enrichment analysis ^56^ presented in Figure 6 was performed using the log_2_ fold change values between the A2780Cis (resistant) and A2780 (parental) cell lines at the protein abundance and protein degradation (*k*_deg_) level using a function provided by the Perseus ^57,58^ platform (v1.6.14.0). The annotation of the gene ontology biological process (GOBP_Direct) was extracted from the DAVID database ^59^. In the “bubble plot” presented in Figure 6, only those categories with at least 10 protein IDs and a enrichment P < 0.01 were visualized and further restricted to the top 25 categories with the lowest P values. The size of the dots was used to reflect the number of proteins in a category.

#### Identification of cisplatin sensitivity-related genes using DepMap

The Dependency map (DepMap) portal ^34^ was used to perform a custom correlation analysis to identify gene expressions associated with cisplatin sensitivity. A Pearson correlation was calculated between the mRNA expression dataset (Batch corrected Expression Public 24Q2) and the cisplatin (CIS-DDP) sensitivity data (IC_50_ based on Sanger GDSC1) using all cell lines available (N = 154); and 414 genes were identified with a significant correlation (P < 0.01, as reported by DepMap) and mapped to the protein abundance and degradation-level data for the analysis as presented in Figure 6.

### Data visualization

The following R packages were used for data visualization: ***ggplot2***, ***ggrepel***, ***ggrastr***, ***LSD***, and ***VennDiagram***. In boxplots, the box in the plot represents the interquartile range (IQR), with the lower and upper edges indicating the first quartile (Q1) and third quartile (Q3), respectively, and the line inside the box marking the median. Whiskers extend to the largest and smallest values within 1.5 times the IQR from the edges of the box. Data points beyond this range are considered outliers and are displayed as individual points. In density/violin plots, the density represents a smoothed estimate of the data distribution, computed using a kernel density estimation (KDE) method; the area under the density curve is equal to 1.

### New Data Visualization Features for Multi-Channel Workflows in Spectronaut v19.3

In addition to the specialized multi-channel workflows and channel-Q-value filtering options, new data visualization features for multi-channel data inspection were made available from Spectronaut v19.3 onward. These include e.g., scoring histograms and channel H/L ratio plots, which provide a detailed overview of scoring weights across channels and the overall H/L ratio distribution across multiple samples and time points, facilitating easy experiment quality control. Additionally, new protein-specific H/L plots can be visualized across runs to assess the quality of individual data points.

## Data availability

The mass spectrometry data and raw output tables as results have been deposited to the ProteomeXchange Consortium via the PRIDE ^60^ partner repository with the following identifiers. The 2-channel standard dilution sample of the A2780 cell line was deposited previously with an identifier **PXD021922.** The HeLa 3-channel data were downloaded from **PXD039578.** The A2780 and A2780Cis total proteome and pSILAC experiment and the pSILAC experiment in the fibroblast cell lines will be available upon manuscript acceptance. The RNA sequencing results were downloaded from the GEO under accession **GSE173201.** The ***KdeggeR*** package will be provided via github.com.

