## Supplementary information for "A Comprehensive and Robust Multiplex-DIA Workflow Profiles Protein Turnover Regulations Associated with Cisplatin Resistance"

### Supplementary tables

**Table S1: Results of the statistical analysis (related to Figure 6).** This table presents the statistical analysis results, including significantly altered genes and proteins between A2780Cis and A2780, as determined by a moderated t-test with Benjamini-Hochberg FDR correction.

**Table S2: Results of the Metascape enrichment analysis (related to Figure 6).** This table summarizes the enrichment analysis performed using Metascape, showing a four-list enrichment analysis of protein IDs that were significantly up- or down-regulated at the protein abundance or degradation levels between A2780Cis and A2780.

**Table S3: Results of the 2D enrichment analysis of GOBP terms (related to Figure 6).** This table provides the results of a 2D enrichment analysis of Gene Ontology Biological Process (GOBP) terms. Log2 fold changes (A2780Cis/A2780) at the protein abundance and degradation levels were used as input for the analysis.

**Table S4: Cisplatin-associated genes identified by DepMap, mapped to proteomic data (related to Figure 6).** This table lists cisplatin-associated genes identified through the DepMap project, mapped to the proteomic data. Statistical significance at the protein abundance or degradation level is indicated.

### Supplementary figures

**Figure S1**

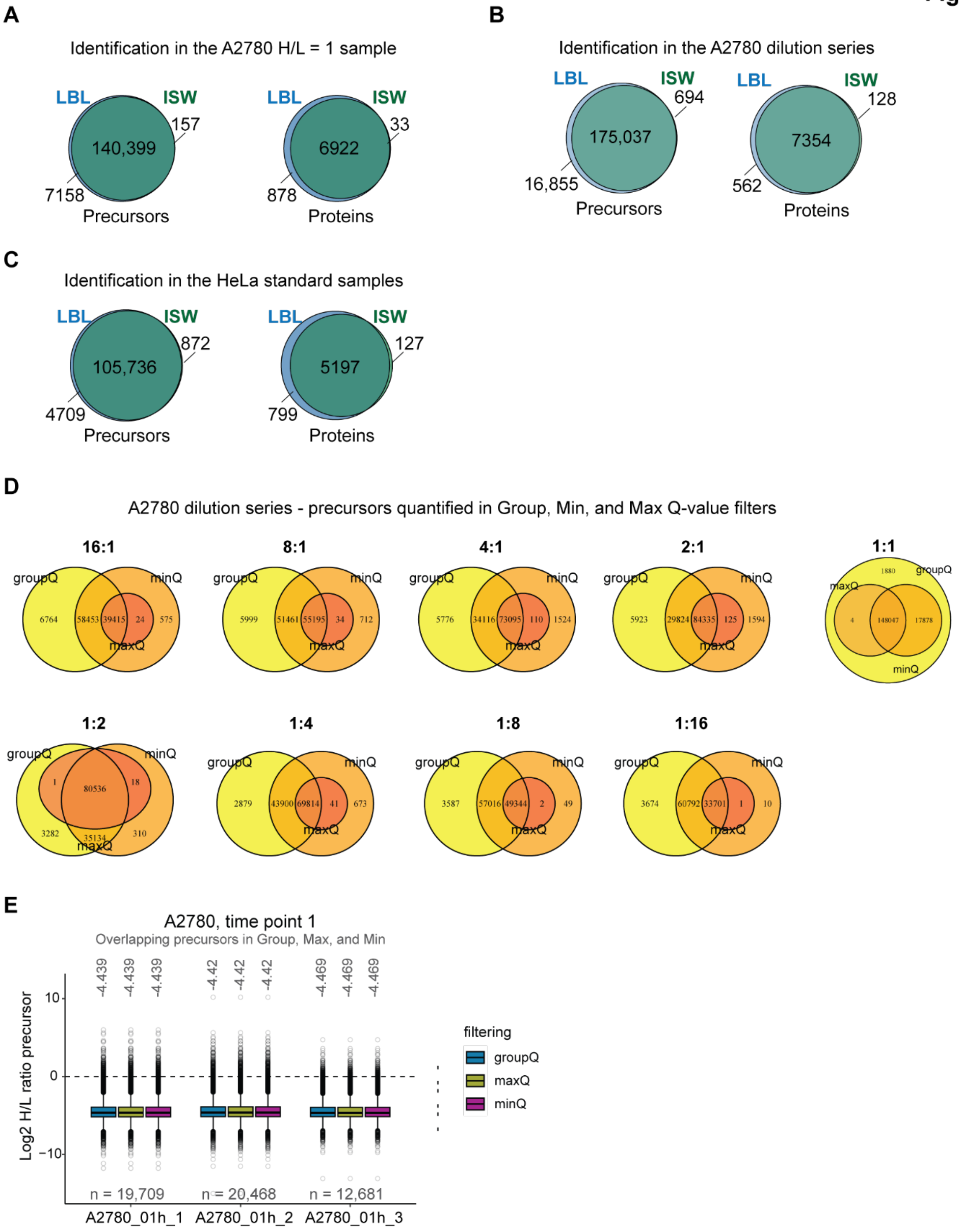


**Figure S1: Identification and quantification using multichannel workflows in Spectronaut (related to Figures 2 and 3): (A)** The number of precursors and proteins in the H/L = 1 A2780

sample identified using the inverted spike-in workflow (ISW) and labeled workflow (LBL) in Spectronaut. **(B)** The number of precursors and proteins in the A2780 dilution series identified using the inverted spike-in workflow (ISW) and labeled workflow (LBL) in Spectronaut. **(C)** The number of precursors and proteins in HeLa standard samples identified using the inverted spike-in workflow (ISW) and labeled workflow (LBL) in Spectronaut. **(D)** The numbers of precursors and their overlap quantified after GroupQ, MinQ, and MaxQ filtering in the A2780 dilution series. **(E)** Precursor H/L ratios in three replicates of the 1-hour time point in the A2780 cell line pSILAC experiment. The precursors were filtered to include only those quantified using all three quantification filtering options in the respective sample. The numbers below the plots indicate the number of overlapping precursors, and the numbers above the plots indicate the medians of the Log2 H/L ratio distributions.

**Figure S2**

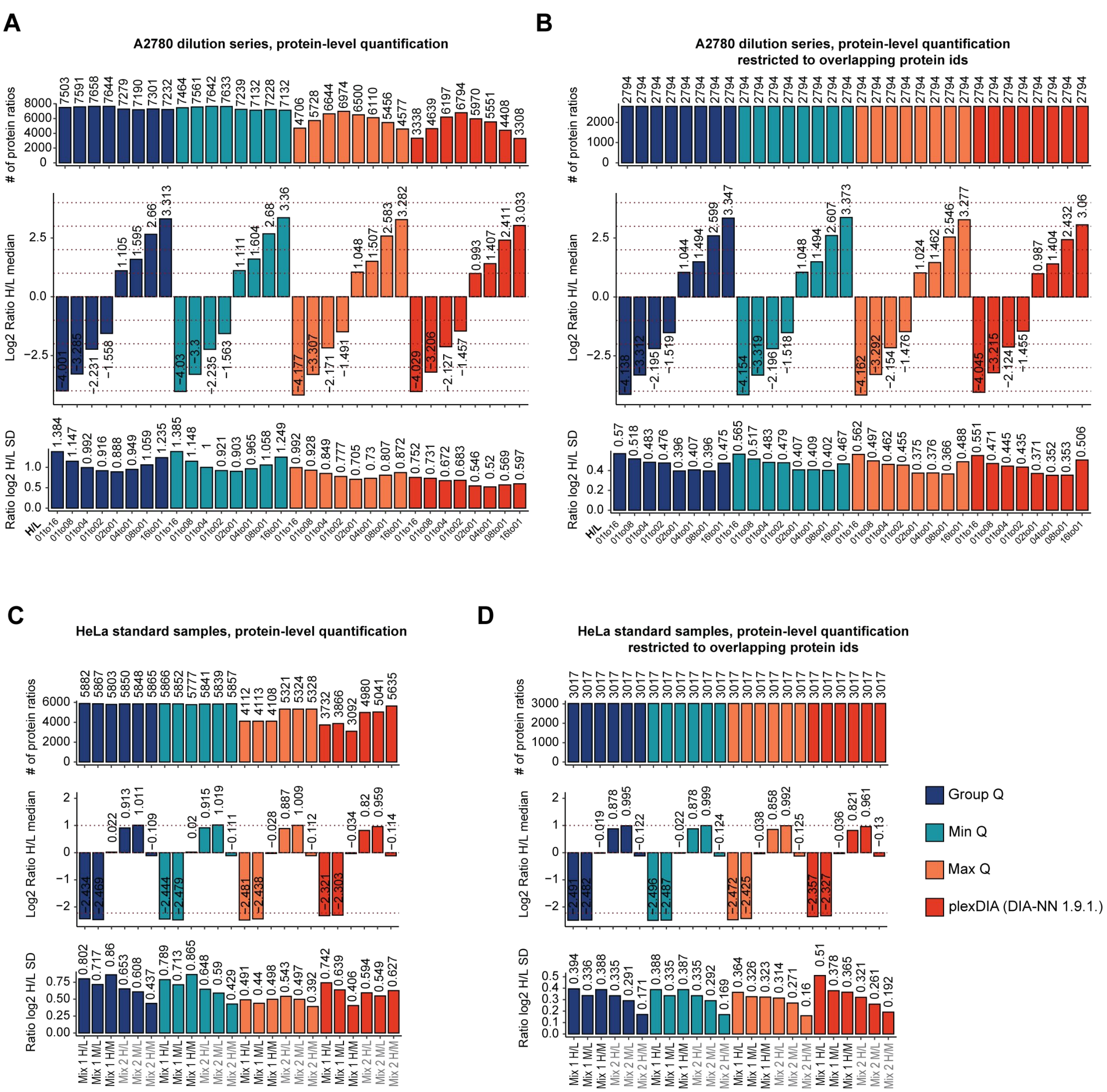


**Figure S2: Quantification of multiplex DIA samples using Spectronaut and DIA-NN (related to Figure 3). (A)** Comparison of the number of quantified protein IDs (upper), the log2 H/L ratio medians (middle), and the log2 H/L ratio standard deviations (lower) in the A2780 dilution series. In **(B)**, the proteins are restricted to a set of protein IDs quantified by both software tools and Q-value filtering strategies. **(C)** Comparison of the number of quantified protein IDs (upper), the log2 H/L ratio medians (middle), and the log2 H/L ratio standard deviations (lower) in HeLa standard samples. In **(D)**, the proteins restricted to a set of protein IDs quantified by both software tools and Q-value filtering strategies. In all plots, data were analyzed using either Spectronaut 19 or DIA-NN v1.9, with additional matrix channel Q-value filtering (Q < 0.01).

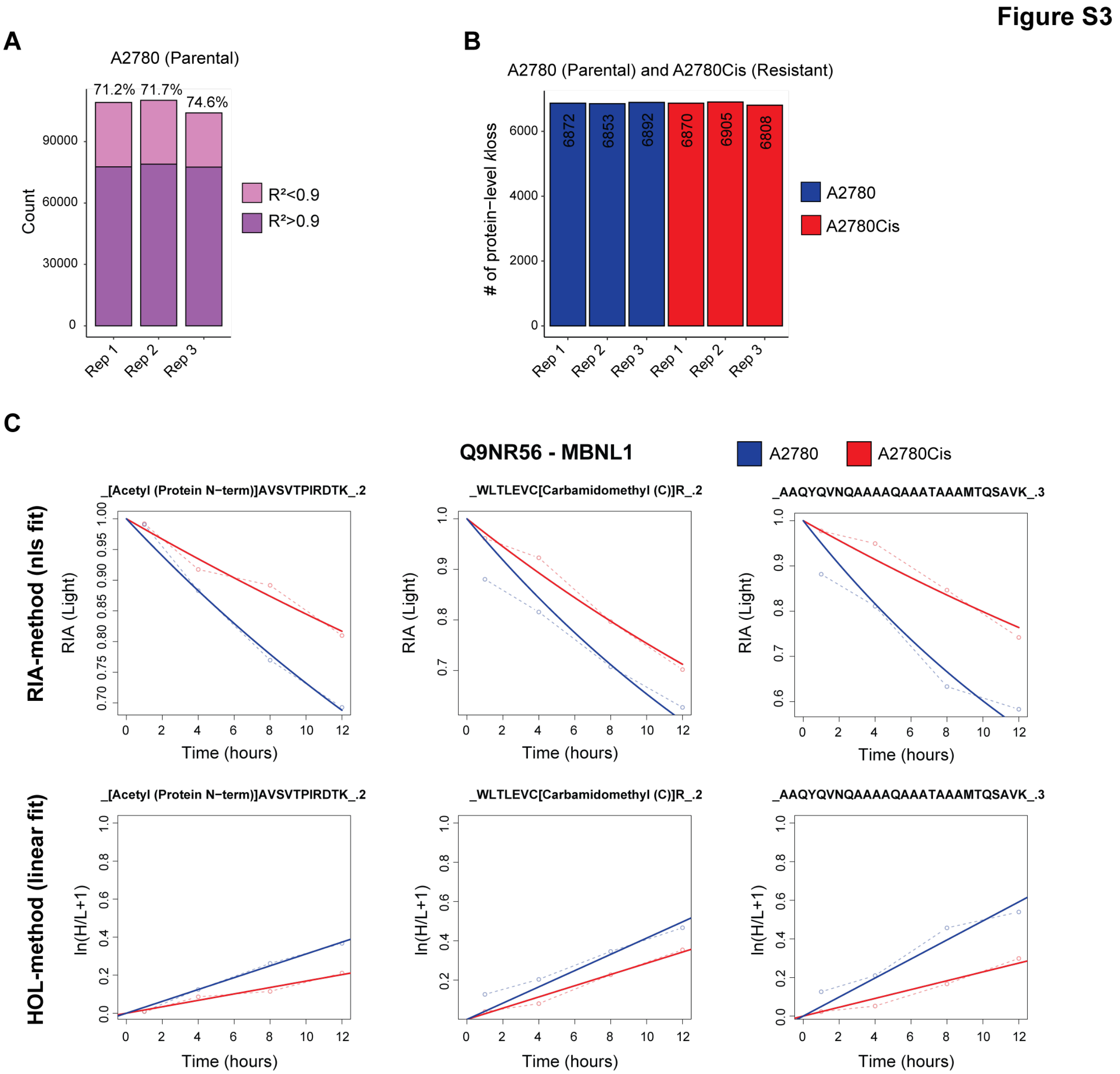


**Figure S3: Protein turnover analysis in the A2780 model (related to Figure 4). (A)** The number of precursors categorized based on the $R^2$ of the linear fit performed using log-transformed H/L ratios and the duration of the pSILAC chase (time point) in three replicates of the A2780 parental cell line. **(B)** The number of protein-level $k_{loss}$ values estimated using KdeggeR in the A2780 and A2780Cis models across replicates. **(C)** Selected examples of the nonlinear least squares (nls) fit using the RIA method and the linear fit using the HOL method. The selected precursors belong to the MBNL1 protein.

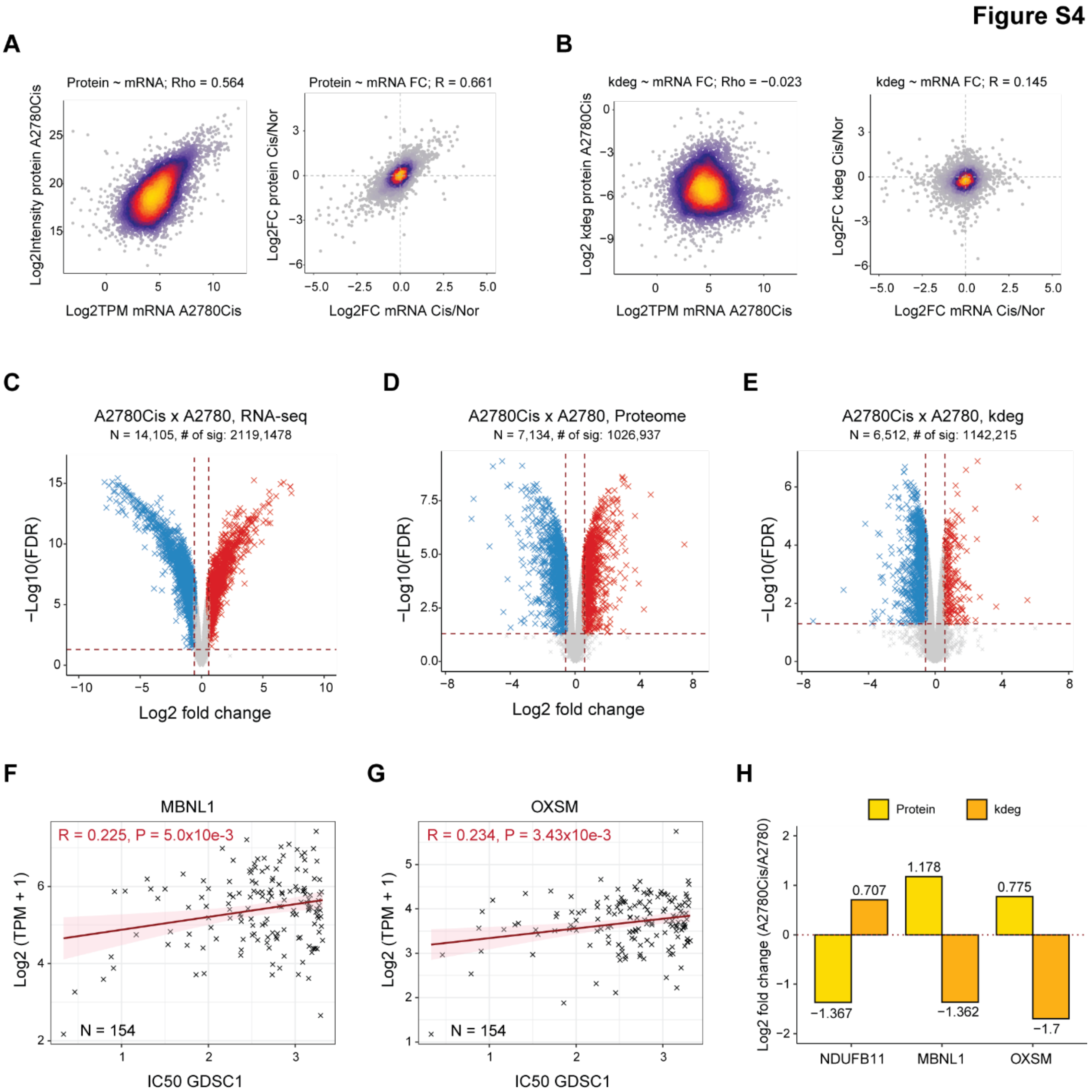


**Figure S4: Multi-omic analysis of the cisplatin-resistant model (related to Figures 5 and 6).** **(A)** Correlation between absolute transcript and protein abundance (left) and relative transcript and protein fold change (A2780Cis/A2780; right). **(B)** Correlation between absolute transcript and protein degradation rates ($k_{deg}$; left) and relative transcript and protein degradation fold change (A2780Cis/A2780; right). **(C-E)** Results of the statistical analysis between A2780Cis and A2780 at the mRNA level **(C)**, protein abundance level **(D)**, and protein degradation level **(E)**. Statistical analysis was performed using a moderated t-test, with the numbers of significantly up- and down-regulated IDs provided (Benjamini-Hochberg FDR < 0.05 and absolute fold change > 1.5). **(F-G)** Positive correlation between MBNL1 **(F)** and OXSM **(G)** gene expression at the mRNA level and cisplatin IC50 based on the GDSC1 dataset, indicating that increased expression of these genes is significantly associated with increased IC50 (resistance). The data were retrieved from DepMap.

**(H)** Histogram of log2 fold changes (A2780Cis/A2780) at the protein abundance and protein degradation levels for NDUFB11, MBNL1, and OXSM.
